# Multi-omic landscape of human gliomas from diagnosis to treatment and recurrence

**DOI:** 10.1101/2025.03.12.642624

**Authors:** Hadeesha Piyadasa, Benjamin Oberlton, Mikaela Ribi, Jolene S. Ranek, Inna Averbukh, Ke Leow, Meelad Amouzgar, Candace C. Liu, Noah F. Greenwald, Erin F. McCaffrey, Rashmi Kumar, Selena Ferrian, Albert G. Tsai, Ferda Filiz, Christine Camacho Fullaway, Marc Bosse, Sricharan Reddy Varra, Alex Kong, Cameron Sowers, Melanie Hayden Gephart, Pablo Nuñez-Perez, EnJun Yang, Mike Travers, Michael J. Schachter, Samantha Liang, Maria R. Santi, Samantha Bucktrout, Pier Federico Gherardini, John Connolly, Kristina Cole, Michael E. Barish, Christine E. Brown, Derek A. Oldridge, Richard R. Drake, Joanna J. Phillips, Hideho Okada, Robert Prins, Sean C. Bendall, Michael Angelo

## Abstract

Gliomas are among the most lethal cancers, with limited treatment options. To uncover hallmarks of therapeutic escape and tumor microenvironment (TME) evolution, we applied spatial proteomics, transcriptomics, and glycomics to 670 lesions from 310 adult and pediatric patients. Single-cell analysis shows high B7H3+ tumor cell prevalence in glioblastoma (GBM) and pleomorphic xanthoastrocytoma (PXA), while most gliomas, including pediatric cases, express targetable tumor antigens in less than 50% of tumor cells, potentially explaining trial failures. Longitudinal samples of isocitrate dehydrogenase (IDH)-mutant gliomas reveal recurrence driven by tumor-immune spatial reorganization, shifting from T-cell and vasculature-associated myeloid cell-enriched niches to microglia and CD206+ macrophage-dominated tumors. Multi-omic integration identified N-glycosylation as the best classifier of grade, while the immune transcriptome best predicted GBM survival. Provided as a community resource, this study opens new avenues for glioma targeting, classification, outcome prediction, and a baseline of TME composition across all stages.

## Introduction

Diffuse glioma are primary brain tumors that are among the most lethal forms of cancer, with limited treatment options and poor prognosis^1,2^. The average survival of patients with glioblastoma (GBM) is only 9–18 months, while diffuse midline glioma (DMG) is the leading cause of brain tumor-related deaths in children^3^. Despite advances in immunotherapies that have shown promise in other cancers, outcomes for gliomas have remained stagnant for decades, and the mechanisms underlying this therapeutic resistance remain elusive^4^.

Due to their location and invasive growth into the normal brain, complete surgical resection of diffuse gliomas is often not possible making them prone to recurrence^5^. Additionally, the glioma tumor microenvironment (TME) is myeloid rich and highly immunosuppressive, rendering them resistant to treatment^6–8^. Clinical therapies currently in trials are attempting to replicate previous successes in other malignancies that use antibodies or engineered T-cells to target tumor-enriched antigens^9^. Indeed, tumor antigen (TA) expression, such as B7H3 and EGFR, represents a promising avenue for directly targeting tumor cells through CAR-T cell therapies and monoclonal antibodies^10–12^. Still, these therapies have either been ineffective or have led to only modest improvements in patient survival^13^. Perhaps related to these setbacks, the expression patterns of these TAs in human lesions are largely unknown^14,15^. Much of what is known about TAs and glioma TME has been gleaned from bulk transcriptomic analysis, patient-derived xenograft mouse models, and tumor cell lines^16–20^. Taken together, these findings point to a significant gap in our understanding of how TAs, immune-tumor interactions, and the functional states induced by them mutually reinforce one another to maintain a treatment resistant milieu. Additionally, though post-translational molecular features, such as tumor glycosylation, have been shown in other malignancies to play a crucial role in promoting tumor-immune-suppression, the glycome in human gliomas remains largely unexplored^21–23^.

To address these questions, we assembled a clinically annotated human glioma tissue cohort from multiple sites across the US. This cohort includes 677 samples from 310 adult and pediatric patients that are classified as high or low-grade tumors, and encompass treatment-naive, post-immunotherapy (anti-PD1, vaccine, anti-CD27, combinatorial), recurrent, and longitudinal cases, making it one of the largest and diverse glioma cohorts assembled^24–27^. To capture multiple layers of molecular regulation, we employed complementary spatial profiling technologies to quantify biomolecule abundance, cellular states and identities (i.e., transcriptome, proteome, and glycome). Multiplexed Ion Beam Imaging (MIBI-TOF) provided high-resolution spatial mapping of 1.2 million cells consisting of neurons, endothelial, tumor and all immune cells (with 18 annotated sub-populations), quantifying targetable tumor antigen expression (i.e., B7H3, EGFR) and immune regulatory receptors (i.e. PD-L1, TIM3) at the single-cell level^28^. Co-registered tissue sections linked these single-cell maps to spatially resolved transcriptomic profiles of over 11,000 genes from immune-cell-rich versus-poor regions using digital spatial profiling (DSP)^29^. Lastly, co-registered tissue sections were also combined with spatial maps of the *N*-glycome quantified by matrix-assisted laser desorption-ionization mass spectrometry imaging (MALDI-MSI) which discovered 70 differentially expressed glycans across gliomas that that could influence immune recognition and tumor behavior^30–32^. Combining these modalities, we established direct correlations between cell composition, protein expression, gene expression, and glycan profiles within the same spatial context across our diverse glioma cohort and how it relates to clinical metadata.

Multi-modal data analysis across these lesions provided new insights into how the glioma TME is shaped by tumor classification and treatment, revealing potential explanations for the limited efficacy of TA-targeted immunotherapies. Furthermore, both tracing the evolving structure of gliomas from primary to recurrent disease and aligning molecular data with patient metrics uncovered factors linked to overall survival^33^. Towards this, we conducted a spatial proteomics-based, quantitative assessment of glioma TAs, marking the first time the prevalence of these antigens and their co-occurrence within single cells has been systematically profiled. Our results indicate that, aside from the high abundance of B7H3+ tumor cells in GBM and pleomorphic xanthoastrocytoma (PXA), most tumors only show less than 50% TA-positive cells. Yet, co-expression analyses suggest that dual-agent strategies could meaningfully increase the fraction of tumor cells potentially targeted. Consistent with the modest clinical improvements seen to date, comparing lesions from patients who received immunotherapy versus standard of care revealed only minor differences tumor architecture^2^. To better understand the apparent stalling of anti-tumor immune activation, we next characterized the glioma glycome, identifying sialylated and core-fucosylated N-glycans whose shifts in abundance correlated strongly with tumor grade, representing both a new classification and therapeutically targetable entity. Spatial transcriptomics further reinforced molecular pathways influencing glycosylation in glioma biology^6^. Still, examining patient outcomes in primary GBM, we found that immune cell, as opposed to tumor cell, programs played the most prominent role in predicting overall survival. All the underlying data from this diverse glial tumor cohort is fully accessible, without restriction, via an interactive online portal (www.bruce.parkerici.org), providing a multi-omic resource to the community that should prove invaluable for future clinical design, beyond the insights provided herein.

## Results

### Diverse and clinically detailed glioma cohort and applied spatial multi-omics

The goal of this study was to create a more comprehensive understanding of the glioma TME and how the multitude of regulatory features at play may integrate to reconcile disease diversity, including the prevalence and amounts of targetable TA. To achieve this, we collaborated with five clinical sites across the US to build a multi-omic atlas of the glioma TME, assembling a cohort of glioma patients with detailed clinical annotations, covering a wide age range from under 1 year to 88 years (**Figure 1A, Figure S1A, Table S1**). The cohort spans World Health Organization (WHO) grades 2, 3, and 4, and includes patients diagnosed with GBM, IDH-mutant astrocytoma, IDH-mutant, oligodendroglioma, IDH-mutant and 1p/19q-codeleted PXA, DMG and other pediatric high-grade gliomas (pHGG). We also included two cohorts of recurrent GBM patients treated with neoadjuvant anti-PD1 immune checkpoint blockade, as well as longitudinal samples from patients with primary and recurrent IDH mutant astrocytoma, oligodendroglioma, and PXA, some of whom received neoadjuvant immunotherapy before surgical resection (**Figure 1B, Figure S1A, Table S1**). Clinical data, including mutational profiles, such as IDH status (Isocitrate Dehydrogenase), MGMT methylation (O6-Methylguanine-DNA Methyltransferase), PTEN (Phosphatase and Tensin Homolog), p53 (Tumor Protein 53), and ATRX status (Alpha Thalassemia/Mental Retardation Syndrome X-linked) along with overall survival, corticosteroid use, and other key attributes, were harmonized across all centers to create a unified, clinically annotated dataset (**Figures 1A and 1B, Figure S1A, Table S1**). This integrated resource enabled us to perform in-depth analyses and explore the molecular intricacies within the TME.

**Figure 1.**
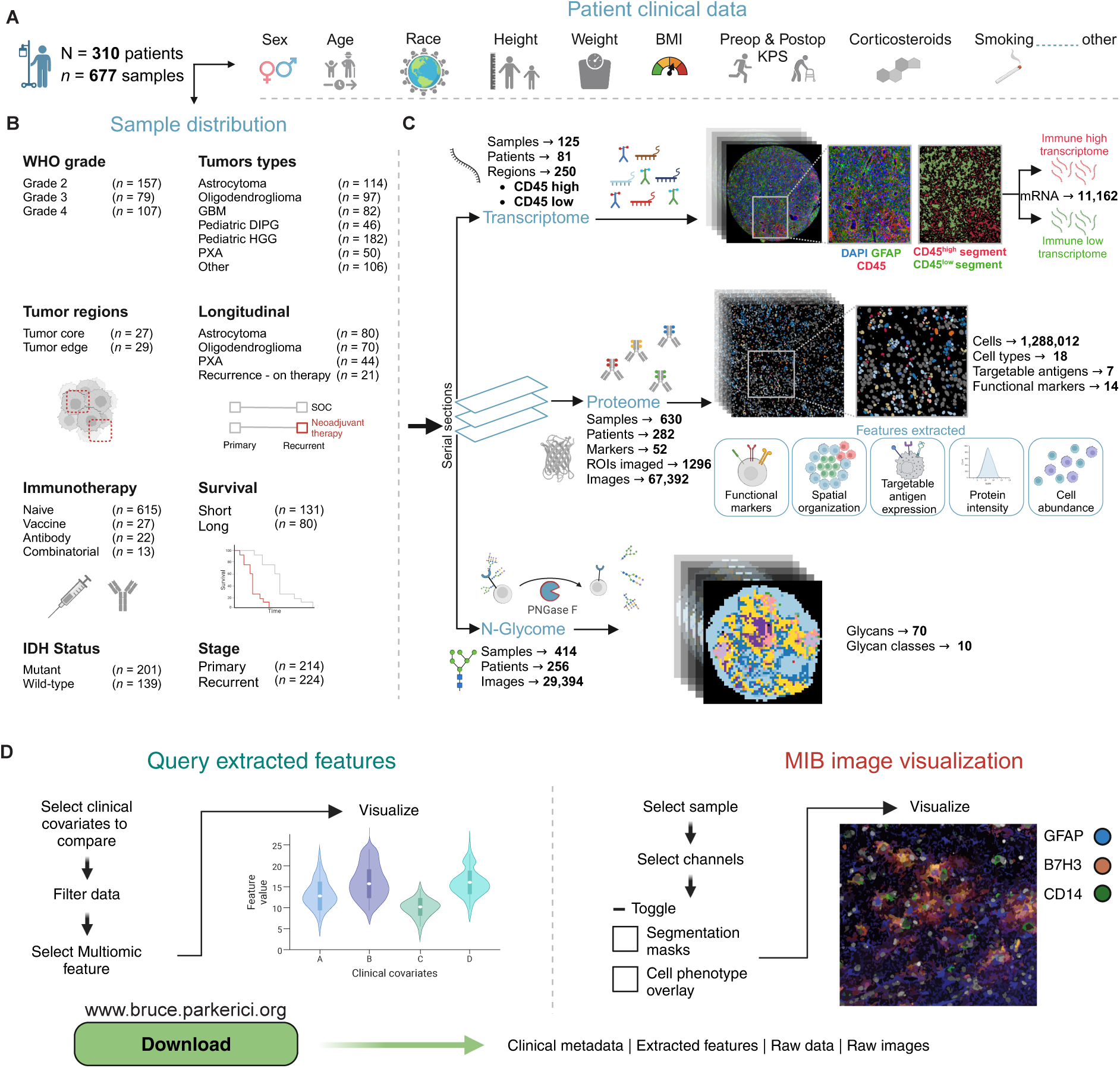
BRUCE (BRain tUmor heterogeneity deCiphEred by high dimensional multi-omic analysis) cohort and data overview. **(A)** BRUCE cohort summary (310 patients, 677 samples), including patient demographics, **(B)** sample distribution and **(C)** the spatial modalities (transcriptomics, proteomics, N-glycome) used for data acquisition from FFPE sections **(D)** Illustration of web resource.

Multiple regions for each lesion were analyzed using single cell spatial proteomics, spatial transcriptomics and spatial N-glycomics using adjacent serial sections to map cellular phenotypes, whole transcriptome in immune-rich versus-poor regions, and N-glycan abundances and distribution (**Figure 1C, Figure S1A**). In total, we imaged over 1.2 million cells from 677 patient samples quantifying the expression and cellular distribution of 52 proteins. The functional status of these cells was characterized using 14 immune regulatory markers and 8 TA across more than 1,000 sampled tumor regions. Our whole transcriptome analysis revealed over 11,000 genes enriched in immune cell-rich and-poor regions, while N-glycomics uncovered 70 unique N-glycans across 10 broad classes, including sialylated structures implicated in immune suppression within the TME of other cancers^34–36^.

We have made all data from this study available on an intuitive, open-access portal, www.bruce.parkerici.org, allowing researchers and clinicians to explore complex spatial and multi-omic data alongside clinical metadata, providing meaningful insights that may guide future research and clinical decisions (**Figure 1D**). The platform, accessible even to non-computational experts, includes interactive tools such as Vitessce for single-cell spatial MIBI image exploration^37^. Additionally, all raw data, including single-cell tables, glycan expression data, spatial glycan TIFF files, and raw MIBI images, are freely available for download without restriction.

### A spatial atlas of glioma: Single-cell profiling

Understanding the composition, distribution, and antigen expression of immune and tumor cells in human gliomas is crucial for understanding drivers of therapeutic failure and for developing new treatments^20,27,38,39^. With this in mind, we used spatial proteomics to provide a basic level of identity to all cells within the TME, including the quantification of tumor cells and 18 distinct immune cell subsets (**Figures 2A and 2C, Figures S2A – S2N, Tables S2 and S3**). Tumor cells were labeled based on histology by expert pathologists and multiplexed phenotype. For the latter, cells were labeled tumor if they were positive for any combination of Olig2, GFAP, CD133, B7H3, EGFR, GM2/GD2, HER2, VISTA, GPC2, NG2, EGFRvIII, H3K27M, or IDH R132H while being negative for immune, endothelial, or neuronal markers (**Figures S2M**). This approach was taken because glioma tumor cells are nondescript and difficult to identify reliably based on a single protein^40^. However, tumors with K27M mutation of the histone 3 isoforms (H3K27M) or R132H mutation of IDH are an exception because mutation-specific antibodies can be used to specifically identify tumor cells^41,42^. We used this subset of lesions to assess the accuracy of multiplexed tumor phenotyping, demonstrating strong concordance (**Figures S3A – S3D**). The immune compartment (18% of total cells) was dominated by multiple subsets of microglia, macrophages, dendritic cells (DCs), and monocytes that were delineated by coexpression of CD11b, HLA-DR, CD14, CD68, CD163, CD206, CD209, TMEM119 and CD141 (**Figure 2B, Figures S2M and S2N**). Lymphoid cells were sparse (<1% of total cells) consisting of mostly CD8+ and CD4+ T cells (**Figure 2B, Figures S2M and S2N**).

**Figure 2.**
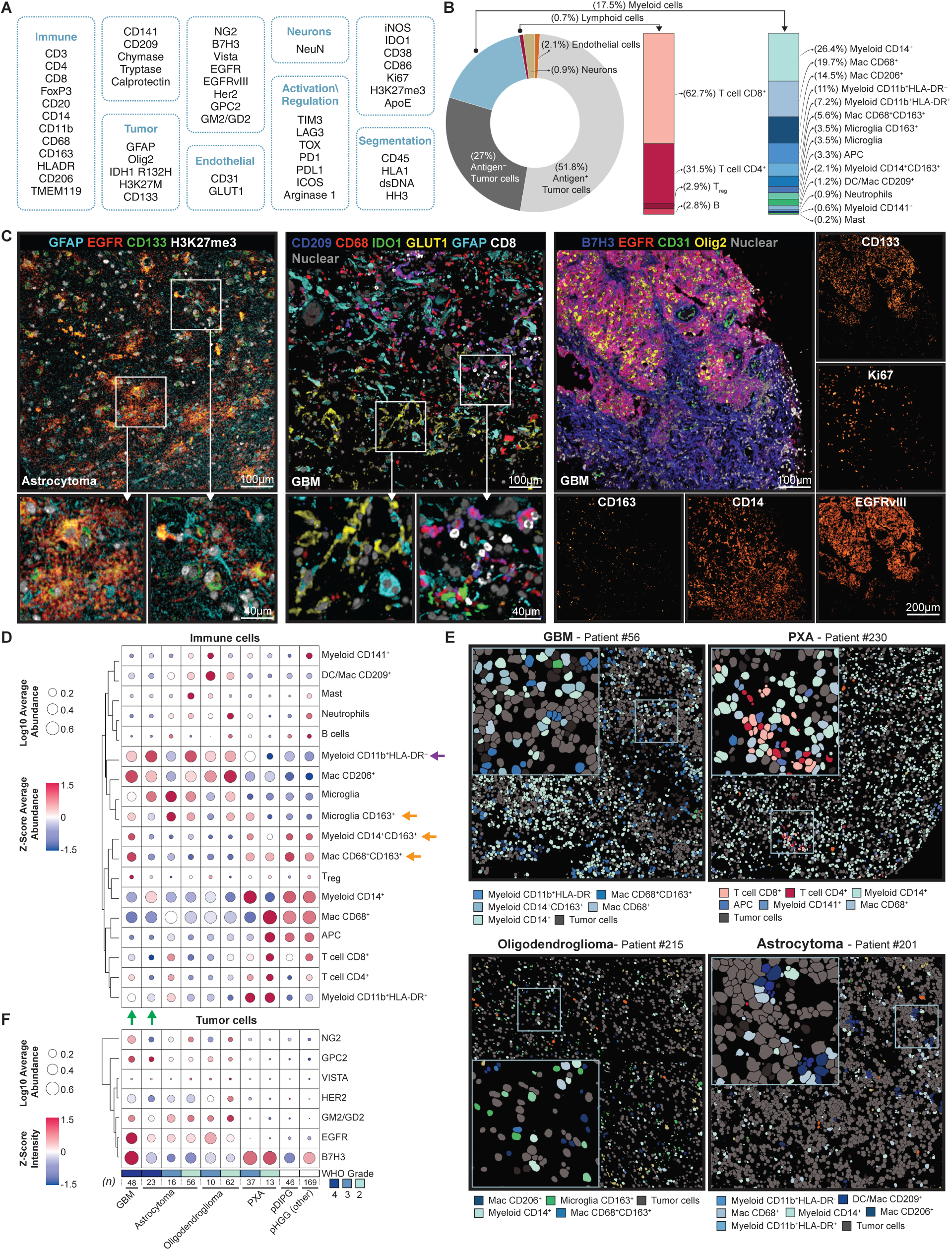
Immune cell and targetable tumor antigen (TA) characterization across glioma subtypes using spatial proteomics. **(A)** Antibody marker panel used to identify various cell types including tumor, immune, neuronal, endothelial, and regulatory cells, as well as segmentation markers. **(B)** Cellular composition of the tumor microenvironment, highlighting targetable antigen positive tumor cells and immune cells, with a detailed breakdown of myeloid and lymphoid subpopulations. **(C)** Representative MIBI images highlighting various markers. **(D)** Heatmap displaying immune cell populations across different glioma subtypes. Color represents row wise Z-score average abundance and circle size represents Log10 average abundance (relative to all immune cells). **(E)** Representative spatial maps showing immune and tumor cell distribution in glioma subtypes, including GBM, PXA, oligodendroglioma, and astrocytoma. **(F)** Heatmap displaying TA across glioma subtypes. Color represents row wise Z-score intensity and circle size represents Log10 average abundance (relative to all immune cells)

Because this cohort spans across glioma type and grade, we were able to identify how cellular composition varies with respect to these clinical covariates (**Figures 2D and 2E, Figure S3E**). For instance, CD163 expression, linked to macrophage wound healing and anti-inflammatory behavior increases with WHO grade and is negatively correlated with overall survival in GBM^43–45^. However, it remains unclear which specific myeloid subsets in each glioma type express CD163^46^. In our cohort, we identified three major myeloid subsets expressing CD163: CD14+, CD68+, and microglia (**Figure 2D, orange arrow**). While CD14+ and CD68+ subsets were found to increase with grade, CD163+ microglia decreased with grade (**Figure 2D, orange arrow, Figures S2F – S2H**), with grade 2 and 3 astrocytoma having the highest relative abundance (4.5% and 7.3% respectively) (**Figure 2D**). Within GBM (grade 4), the proportion of CD14+ and CD68+ subsets expressing CD163 was remarkably high (15% and 29%) (**Figure 2D**). Interestingly, IDH1-mutant grade 4 astrocytoma exhibited a lower abundance of CD163+ myeloid populations compared to GBM, despite containing relatively high levels of total microglia, CD14+, and CD68+ myeloid cells (**Figure 2D, green arrow**). These findings highlight the nuanced CD163+ immune subsets across gliomas and grades and offer new insights into their potential roles in the TME.

These differential immune compositions measured also extend to rare gliomas, such as PXA, an astrocytic tumor affecting children and young adults accounting for less than 1% of gliomas. Due to the rarity of PXAs, our understanding of their TME remains limited^47^. To address this, we characterized the immune cell composition and TA expression across 15 pediatric (Pediatric HGG [other]), 13 grade 2, and 37 grade 3 PXA samples (**Figure 2D and 2E**), which revealed two notable insights. First, PXAs had the lowest relative abundance of myeloid cells phenotypically consistent with MDSCs (CD11b+, HLA-DR-) (**Figure 2D, purple arrow**). Second, Grade 2 PXAs had the highest abundance of T cells across all gliomas. Together, these findings highlight how the immune landscape can remodel with glioma type and grade, implicating the likely need for more nuanced immune targeting strategies instead of a one-size-fits all in glioma.

### Quantifying tumor antigen variability: Implications for targeted therapy in gliomas

Beyond immune cell modulation, targeting of TAs remains a significant strategy for gliomas, in general. Still, the efficacy of TA-targeting therapies is dependent, in large part, on abundant and high frequency expression across tumor cells^48^. However, quantitative profiles describing how these antigens are co-expressed at single cell level with respect to tumor structure, type, and grade in human gliomas are not known. To this end, we quantified the expression of eight different targetable TA (B7H3, EGFR, EGFRvIII, HER2, VISTA, GPC2, NG2, GM2/GD2) that are currently in pre-clinical or clinical stage of investigation in glial tumors^49–56^. EGFRvIII was excluded from analysis due to the limited number of confirmed patients in our cohort with this variant. Because lipids like GD2 are typically lost during processing of archival tissues, we used GD2 synthase (GM2/GD2) as a surrogate for this TA. Of the TAs measured here, B7H3 showed the highest abundance across all tumor types and grades (**Figure 2F**). However, B7H3 was not restricted to tumor cells, with >60% immune and endothelial cells expressing this as well (**Figure S3I**). EGFR was also prominent but only in adult tumors (**Figure 2F**). We further investigated the correlation between the expression of these antigens and their transcripts using spatial transcriptomic data from serial samples. B7H3 and EGFR proteins showed significant correlation with their respective transcripts, while other targetable tumor antigens showed no significant correlations (**Figure S3J**).

We next examined *in silico* how tumor targeting might be improved using multi-agent therapy^57,58^. In high-diversity tumors (GBM, astrocytoma, oligodendroglioma) that more frequently express multiple TAs (**Figure 3A**), our analysis suggests that dual therapy could lead to a significant improvement in tumor coverage–defined here as the percentage of tumor cells expressing at least one of the TAs of interest (**Figure 3B**). However, this benefit quickly plateaus and does not improve further with three or more agents. In contrast, because low-diversity tumors (**Figure 3A**) mostly expressed B7H3, little improvement in tumor coverage was seen by targeting multiple TAs (**Figure 3C**). Given that dual-TA coverage was most effective in high-diversity tumors, we assessed TA pairings to determine the optimal combination for each patient (**Figure 3D, Figure S3K**). Dual therapy with B7H3 and EGFR was predicted to be optimal, covering up to 100% of tumor cells, for 47% of GBM and 44% of oligodendroglioma cases. In astrocytoma, no dominant pair emerged, though 22% of patients had either B7H3 with EGFR or HER2 (**Figure 3D**).

**Figure 3.**
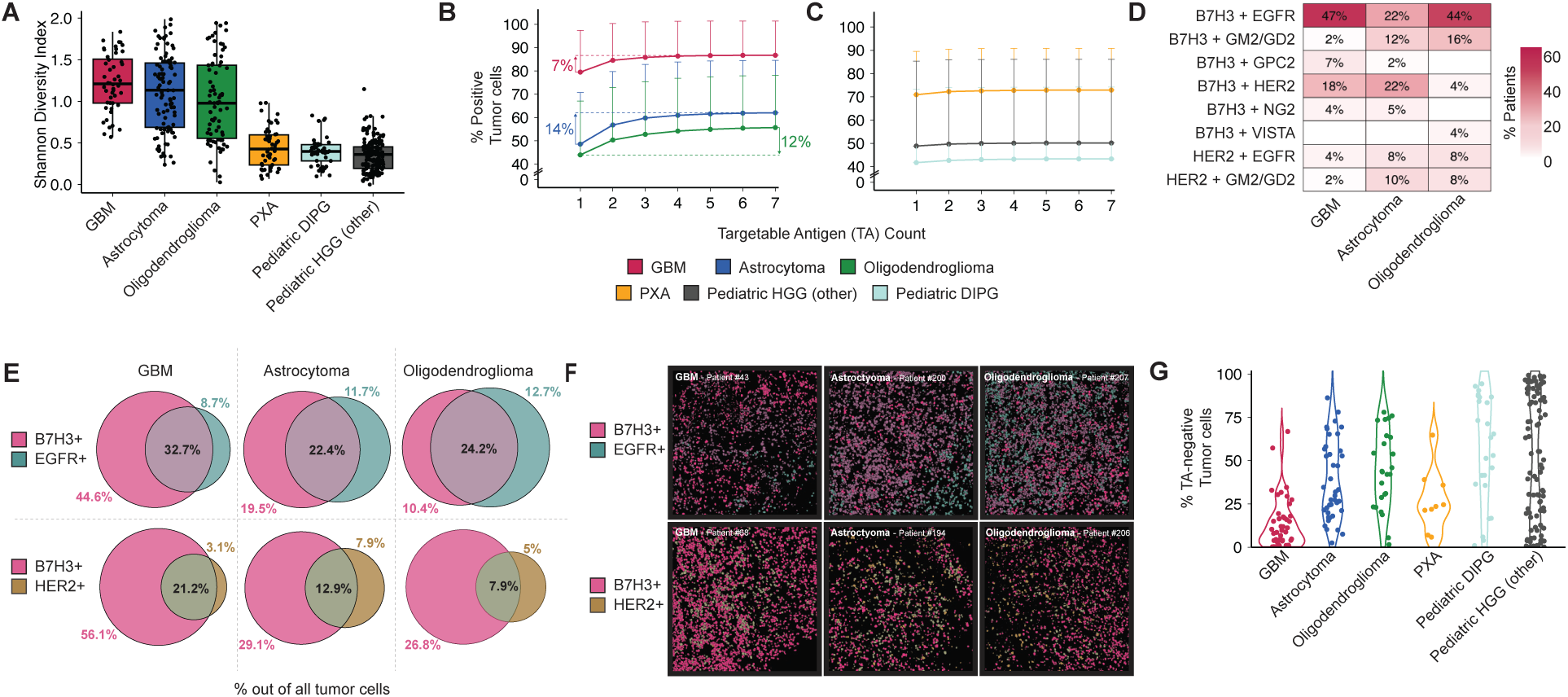
Comprehensive analysis of tumor antigen diversity, coverage, and coexpression in gliomas. **(A)** Boxplot of Shannon diversity index showing TA heterogeneity across different tumor subtypes. **(b-c)** Line graph showing % tumor cell coverage with increasing TA. Error bars show SD. **(D)** Top two tumor antigen expression heatmap shows % patients for combinations of selected top 2 TA yielding maximal coverage of tumor cells for high TA diversity glioma. **(E)** Venn diagrams showing % of tumor cells positive and overlap for B7H3, EGFR and HER2 out of total tumor cells in high TA diversity gliomas. **(F)** Representative tumor CPM images, illustrating the distribution of tumor antigens for coverage and coexpression. **(G)** Violin plot of % tumor cells negative for the 7 TA measured. Each dot represents an individual patient.

Convoluting this, antigen expression by non-tumor cells, as we saw with B7H3, can lead to off-target effects and adverse events when using monospecific therapies alone or with multi-agent regimens like the ones described above^59^. To circumvent these issues, bispecific antibodies and CAR-T cells that require co-expression of two antigens on the same cell are being examined as a means of improving tumor specificity^57,58^. With this in mind, we quantified how often B7H3, EGFR, and HER2 were co-expressed on the same cell in high-diversity tumors to understand how tumor coverage with a bispecific therapy compared with monospecific dual therapy (**Figures 3E and 3F**). Overall, we found the added specificity of a bispecific therapy is predicted to come at the cost of reduced tumor coverage. For example, in GBM, 85% of tumor cells express either B7H3 or EGFR, but only 32% expressed both. Lastly, we would like to note that nearly all lesions contained tumor cells that were pan-negative for all TAs quantified here, with their prevalence varying widely from 0-99% (**Figure 3G**). Taken together, these results provide new guidance on how therapeutic regimens might be improved while also highlighting the challenges in designing broadly reactive, specific therapies in glioma.

### Benchmarking tumor antigen expression dynamics for targeted therapy in gliomas

Besides frequency, the ideal therapeutic target would be consistently expressed by all tumor cells, across patients, minimizing cell-to-cell variability within each patient’s lesion^57–59^. Given the prevalence of pan-negative tumor cells (**Figure 3G**), we next examined this more broadly to understand how the frequency of TA-positive cells could impact the potential efficacy of tumor cell targeting therapies. To contextualize these findings, we first compared our data with a tumor-targeted therapy that has shown durable survival benefits, CD19 CAR-T therapy in B-cell malignancies. Previous studies have shown a cohort median of 98% antigen-positive tumor cells with individual patients ranging from 72%-100%^60^. Since B7H3 and EGFR emerged as the top TAs for GBM, oligodendroglioma, and astrocytoma, we compared the percentage of tumor-positive cells for these antigens with the rate of CD19 positivity in B-cell malignancies. This analysis revealed two insights. First, of the three tumor types, GBM had the highest fraction of patients falling in this range for both B7H3 and EGFR (62% and 30%, **Figures 4A and B**, respectively). Second, and perhaps encouragingly, TA-positive cells for lesions falling within this range also expressed on average higher amounts of antigen than those outside of it (B7H3 = 1.6X higher, *p = 0.001* and EGFR = 4.2X higher, *p = 0.007*). However, cell-to-cell variability in TA expression (i.e., intra-patient variation) was also greatest in these lesions. Given these findings, we then examined how TA expression varied between patients. We quantified the dynamic range of TA expression (5^th^-95^th^ percentile) at a single-cell level across all tumor cells for all patients (**Figures 4C and 4D**). Similar to our findings when examining variation within a single lesion, we found EGFR and B7H3 to exhibit the largest interpatient variation as well, spanning a dynamic range of 3.49 and 2.59 Log2 FC from median, respectively.

**Figure 4.**
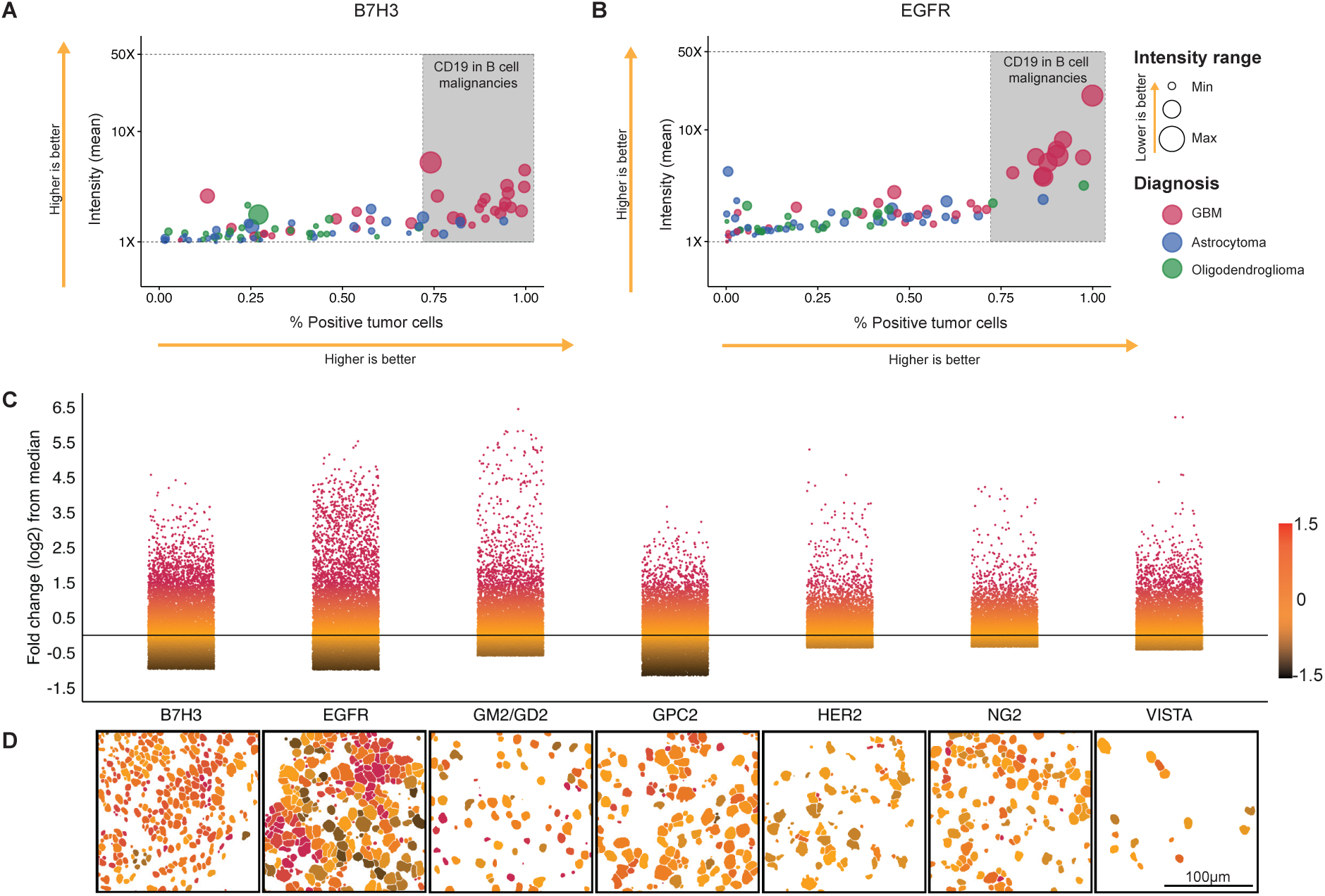
Dynamic range of expression of targetable tumor antigens (TA) in gliomas. Scatter plot for **(A)** B7H3 and **(B)** EGFR showing mean intensity on Y axis, and % positive tumor cells on the x axis. Size of each dot indicates the intrapatient dynamic range of expression. Each dot represents an individual patient. Colors identify tumor type. Shaded region shows theoretical mean intensity and % positive tumor cells of CD19 in B cell malignancies. **(C)** Point plot showing the log2 fold change from the median intensity. 10,000 cells per TA sampled. Color scale represents Log2 fold change capped at-1.5 and 1.5 **(D)** Representative MIBI CPM of tumor antigen expression. Color scale shows the log2 fold change from median shown in panel a. Scale bar = 100 μm.

TA expression is often not spatially uniform throughout and can be biased to the central tumor core or peripheral edge near the resection margin where residual disease is likely to occur^19,61^. With this in mind, we further examined regional variability of TA expression in WHO grade 4 tumors, focusing on the gradient from core to infiltrating edge (**Figures S4A – S4H)**. Here, we found that two of the TAs exhibited non-uniformity with respect to these clinically significant histologic regions. GM2/GD2 tended to be higher at the resection margin while B7H3 was typically higher in the tumor core (**Figures S4A – S4H**). Taken together, these findings underscore the complexity of developing effective therapies for gliomas, as TA variability, both within and between tumors, complicates consistent targeting, even for abundant antigens like B7H3 and EGFR.

### Clinically significant changes in disease status are spatially encoded in the glioma TME

In previous work examining carcinomas, we found clinically significant events like disease recurrence and metastasis to be tightly linked to corresponding changes in TME structure that revealed insights into drivers of tumor progression and therapeutic resistance^8,62–65^. To determine if TME structure and disease progression are similarly linked in gliomas, we compared longitudinally matched primary and recurrent lesions from patients with LGG (Astrocytoma IDH-mutant, Oligodendroglioma IDH-mutant 1p19q co-deletion and PXA) that received standard of care (SOC) using QUantitative InterCellular nicHe Enrichment (QUICHE) (**Figures 5A and 5B, green line**)^63^. QUICHE combines graph neighborhood analysis with statistical modeling to identify cellular niches that are differentially enriched in one condition over another. This analysis revealed systematic shifts in how tumor and immune cells colocalize in recurrent LGG disease (**Figures 5C and 5D, Figure S5A**). For example, in primary tumors, there were complex interactions between CD4+ and CD8+ T cells with CD68+CD163+ macrophages, CD14+ myeloid cells, tumor cells and endothelial cells (indicated by interconnected lines while thickness of lines indicate the prevenance across patients) which were absent in recurrent tumors. Instead, microglia with or without CD163, CD206+ macrophages and CD11+HLADR-myeloid cells formed distinct spatial niches with tumor cells in recurrent tumors.

**Figure 5:**
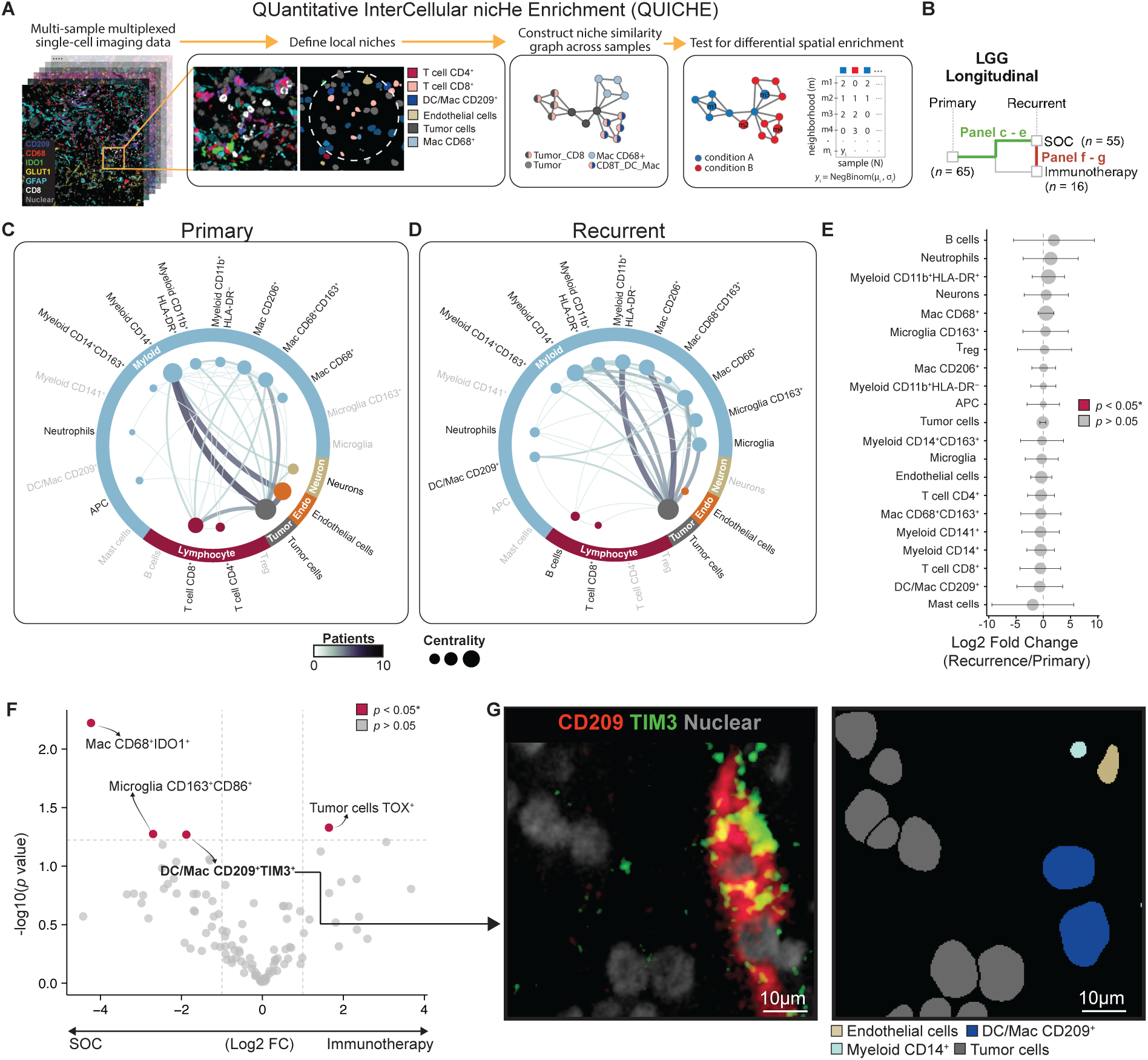
Analysis of distinct spatial niches and therapy-induced changes within the TME of longitudinally sampled low-grade gliomas. **(A)** Schematic overview of the QUICHE analysis pipeline. **(B)** Overview of the longitudinal LGG cohort with paired primary and recurrent samples, including a subset of patients treated with immunotherapy. Spatial interaction graphs derived from differentially abundant niches for **(C)** primary and **(D)** recurrent tumors. Edge width and color corresponds to the number of unique patients with the interaction. **(E)** Differential abundance analysis of immune cell types between recurrent and primary tumors in the SOC cohort, showing log2 fold changes and statistical significance (circle size indicates-log10 p-value). **(F)** Volcano plot comparing log2 fold changes of immune cells with functional marker classification between SOC and immunotherapy-treated patients, highlighting (red) significant differences. **(G)** Representative images of multiplexed single-cell spatial TME landscapes, showing functional marker co-expression within immune cell subsets.

These changes suggest that tumor cells in primary tumors interact more with vasculature and immature myeloid cells in comparison to recurrent tumors where tumor cells interact more with brain resident microglia and more effector myeloid populations^66^. Notably, while the TME structure differed significantly in primary and recurrent lesions, the prevalence of tumor and immune cell subsets composing the TME did not (**Figure 5E**, p > 0.05 for all populations). We next evaluated whether the cells within these differential niches were phenotypically distinct from cells outside of them to understand how TME structure and function are related. TIM3, IDO1, and CD86-positive cell fractions were notably enriched in DE niche neighborhoods (**Figures S5B – S5D**). For example, TIM3+ CD4 and CD8 T cells were overrepresented in primary tumor niches but were entirely absent in recurrent tumor niches. Collectively, these findings demonstrate that although the overall cellular makeup remains constant, glioma recurrence is encoded by the spatial a reorganization of tumor–immune cell interactions.

In addition to patients receiving SOC, our LGG cohort included recurrent samples from phase 1 clinical trials in which patients with Astrocytoma IDH-mutant or Oligodendroglioma IDH-mutant 1p19q co-deletion were treated with either a vaccine alone or a vaccine in combination with an anti-CD27 agonistic antibody^25,26^. Although small sample sizes and limited follow-up precluded a definitive assessment of clinical benefit, immunotherapies in glioma generally have not improved clinical outcome. Thus, we hypothesized that comparison between patients receiving immunotherapy and SOC would not reveal significant spatial differences. In line with this, QUICHE analysis did not reveal any treatment-specific niches. However, bulk frequencies of IDO1+, CD86+, and TIM3+ myeloid cells was higher in SOC-treated patients (**Figures 5F and 5G, Figure S5E**). Taken together, these findings suggest that TME niche enrichment is linked to glioma clinical status through a sensitive and specific encoding that cannot be reliably inferred in a spatially agnostic manner based on cell composition alone.

### Multi-omic profiling of the TME reveals unique glycan profiles across glioma grades and novel cellular and gene module associations

*N*-glycosylation is a post-translational modification occurring on most membrane-bound proteins that regulates a broad range of cell-cell interactions involved in cancer progression and immune regulation^21–23^. For example, tumor sialylated *N*-glycans inhibit innate and adaptive responses through binding of Siglec receptors on immune cells^34–36,67^. With this in mind, we used MALDI-IMS (Bruker TimsTOF MALDI2) to map the spatial composition of glioma TME glycans to determine which structural classes are present and how they change with grade^32,68^. Searching a possible library of N-glycans between 800 and 4000 m/z, we identified 70 unique *N*-glycans across 10 different structural classes (each unique glycan may belong to more than a single structural class) in WHO grade 2-4 tumor samples (**Figure 6A**, **Figure S6A**). The abundance of these glycan classes was grade-specific^69^. For example, grade 2 lesions where enriched in high mannose, fucosylated, hybrid, agalactosylated, and tetrantennary structures while grade 4 gliomas had higher levels of sialylated, polylacnac, bi-and triantennary glycans (**Figure 6B and 6C, Figure S6A**). Notably, we saw that IDH mutant grade 4 tumors were distinctly different from IDH mutant grade 2 and 3 tumors showing lower fucosylation and higher sialylation similar to GBM (**Figure S6B**) indicating that the glycocalyx may be independent of IDH status.

**Figure 6.**
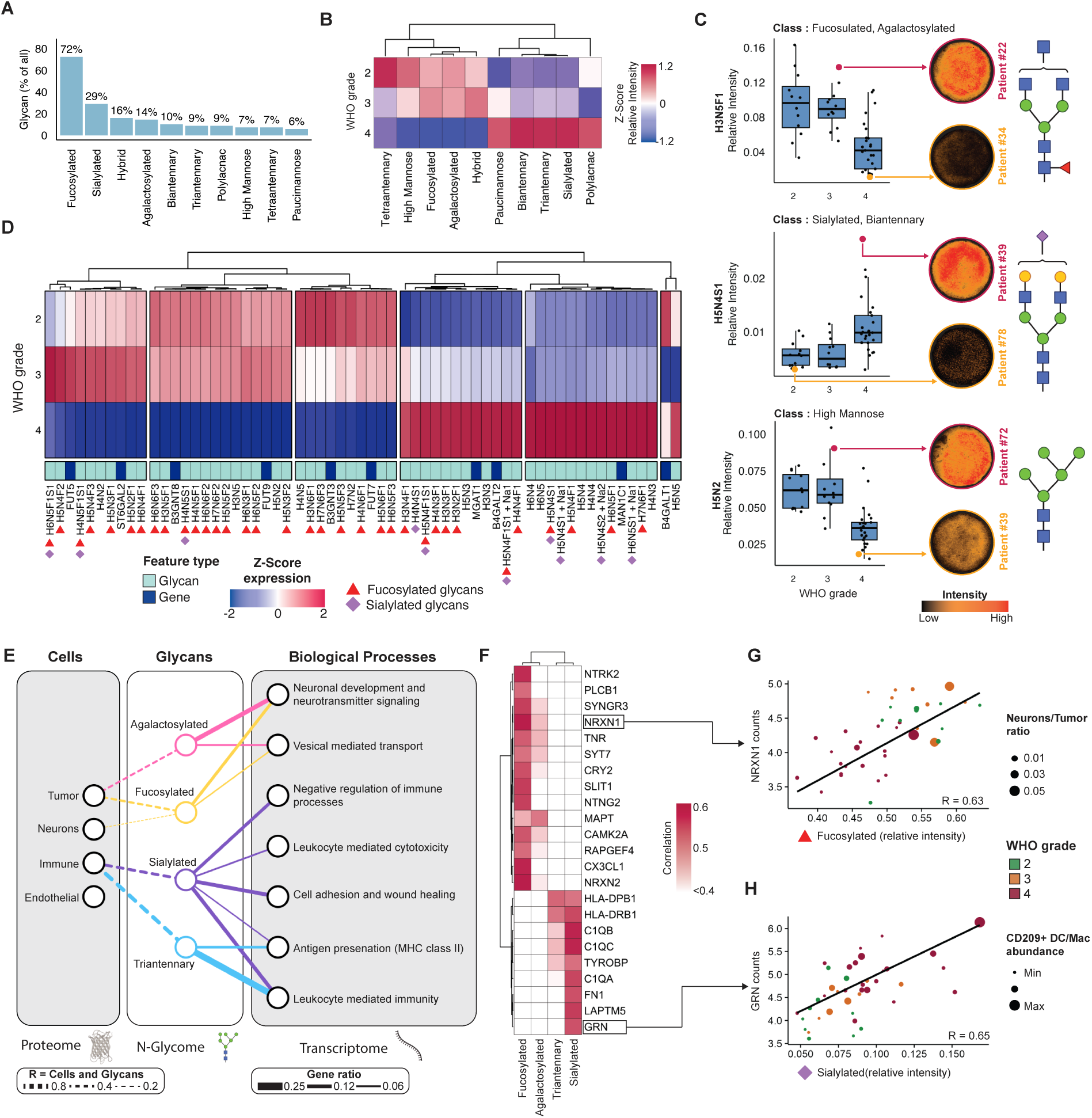
Multi-omic profiling of the TME. **(A)** Bar graph % of glycans identified in each N-glycan class across glioma samples **(B)** Heatmap depicting the relative intensity (Z-score) of glycan classes across WHO grades 2, 3, and 4 **(C)** Boxplots of relative intensity for selected glycans from different glycan classes across WHO grades, with corresponding representative images of glycan staining illustrating low and high glycan expression in different patient samples. Glycan structures are displayed beside the boxplots **(D)** Heatmap showing the Z-score expression of both glycans and RNA for glycan processing enzymes that were differentially expressed between any WHO grades. Glycans are identified by cyan, and RNA is identified by dark blue. **(E)** Network diagram showing cell types linked to glycan classes (fucosylated, agalactosylated, sialylated, tri-antennary) and their associated biological processes, derived from multi-omic analysis. Edge weights between cells and glycans indicate correlation coefficient while edge weights between glycans and biological processes indicate gene ratio (genes identified with high correlation / total genes in pathway) **(F)** Heatmap of the top 25 genes correlated with glycan expression, with rows representing glycan classes and columns representing genes, colored by correlation strength. **(G)** Scatter plot illustrating the relationship between NRXN1 expression and fucosylated glycans, with dot size representing the neuron-to-tumor cell ratio and color indicating WHO grade. **(H)** Scatter plot showing the association of GRN expression with sialylated glycans, with dot size reflecting CD209+ DC/Mac abundance and color corresponding to WHO grade.

To understand how the glycocalyx of gliomas could be remodeled with grade, we then integrated differential N-glycan compositions with tissue whole transcriptome profiling (Nanostring DSP GeoMx) from matching regions from adjacent serial sections. Here we identified glyco-regulatory enzymes whose expression correlated with these trends across grade (**Figure 6D**). Of the ten glycan classes analyzed, fucosylated and sialylated glycans exhibited the greatest concordance with the transcript levels of the enzymes predicted to synthesize them (**Figure 6D**, *red triangle* and *purple diamond*, respectively; **Figures S6C – S6D**). Interestingly however, robust trends with gene expression matching N-glycan modifications, like this, were not found for other glycan classes despite19 genes captures by whole transcriptome profiling. This discordance suggests that the stark remodeling in tumor glycans classes with grade cannot be reliably inferred from upstream molecular information. Consequently, the regulatory role of glycans in glioma biology could be a promising, yet understudied, target for both tumor classification and therapeutic targeting.

In order to better approximate the functional role of N-glycans in gliomas we looked to further integrate the differential signals we overserved with the broader set cellular features mapped for the TME. Therefore, we examined associations with both cellular abundances and total gene (>11000 genes) expression to glycan classes. We further identified all transcripts correlated (R² > 0.4) with each glycan class and performed Gene Ontology (GO) analysis (**Figure 6E**). We found that fucosylated and agalactosylated glycans, associated with brain-specific functions, including neuronal development processes, were positively correlated with neuronal and tumor cells and negatively correlated with most immune cell subsets (**Figure 6E**, **Figure S6E**). In contrast, sialylated and tri-antennary glycans, associated with immune-related gene-expression modules, such as antigen presentation and leukocyte activation, showed strong positive correlations with immune cell abundance (**Figure 6E**). These data suggests that distinct glycosylation patterns may play a key role in tumor related biological processes and influence disease state.

Finally, to identify the main drivers of these biological processes, we extracted the top 25 genes based on correlative magnitude with glycan expression (**Figure 6F**). Among these were NRXN1, NRXN2, NTRK2, LAPTM5, and GRN, all recognized for their involvement in glial tumor biology, highlighting their potential roles in the TME^70–74^. In particular, NRXN1 (neurexin 1), a cell surface receptor that binds neuroligins at synapses in the central nervous system, is known to promote tumor growth through excitatory neuronal activity. Its strong positive correlation with fucosylated glycans along with its positive correlation with the neuron-to-tumor cell ratio in patients, highlights a potential link of surface fucose moieties with neuronal activation (**Figure 6G**). In contrast, GRN (granulin) showed a strong correlation with sialylated glycans. GRN has been indicated to play a role in glioma tumor cell proliferation with increasing expression seen with tumor grade^73,74^. Interestingly, our multi-omic analysis revealed that GRN showed a higher weighted correlation when considering the abundance of CD209+ DC/Mac (**Figure 6H**). This correlation suggests a potential involvement of specific myeloid subtypes with increased sialylation involved in tumor growth. Together, these findings reveal a cohesive picture of glycan involvement within the TME that was reinforced by correlations with cellular abundance and GO-enriched biological pathways. This consistency across three orthogonal modalities provides a holistic view of how glioma glycosylation could impact clinical outcomes.

### Multi-omic integration identifies distinct grade-and survival-specific TME features in GBM

Integrating cellular protein, tissue glycan, and lineage gene expression can provide a deeper understanding of the tumor microenvironment by capturing the complex interplay between molecular pathways that drive tumor progression and immune responses^8^. Therefore, to understand more holistically which aspects of this regulatory hierarchy drive glioma biology between disease grades we trained a random forest classifier on multi-omic data from primary lesions to predict WHO grade (**Figure 7A**). The model incorporated spatial proteomics (cell abundances, ratios, marker intensity, spatial organization, etc.), glycomics, and transcriptomics from immune-enriched (CD45+) or immune-poor (CD45-) glioma tissue regions. After removing sparse and highly correlated features to reduce noise, the model achieved an AUC of 0.80 across 10 random trials with a permuted baseline of 0.57 (**Figure 7B**). The top 75 features by feature class revealed that glycan features had the highest median importance, followed by MIBI features, RNA from immune-poor tumor regions and RNA from immune-rich regions (**Figure 7C**). Among the most important individual features were CD31 intensity in endothelial cells enriched in grade 4 (**Figures 7D – 7F**); H6N3F1 and H5N5F2, two fucosylated glycans enriched in LGG (**Figure 7D**, **Figure 6D**); and Vascular endothelial growth factor A (VEGFA), linked to vascular proliferation also enriched in grade 4 (**Figure 7G**). Notably, only 10% of the top 75 features were transcripts from immune-rich regions (**Table S4**) suggesting that the transcriptome of immune cells is more stable across grade than tumor cells. Interestingly these patterns were largely maintained when considering IDH mutation status (**Figure S7A**) suggesting that grade 4 IDH mutant astrocytoma is distinct from grade 2 and grade 3 IDH mutant tumors. Overall, in line with the initial N-glycan analysis findings (Figure 6), it appears that grade-driven alterations in glycan composition are key determinants of glioma biology, implicating the tumor glycocalyx as a central orchestrator of the molecular and cellular interactions that drive disease biology.

**Figure 7.**
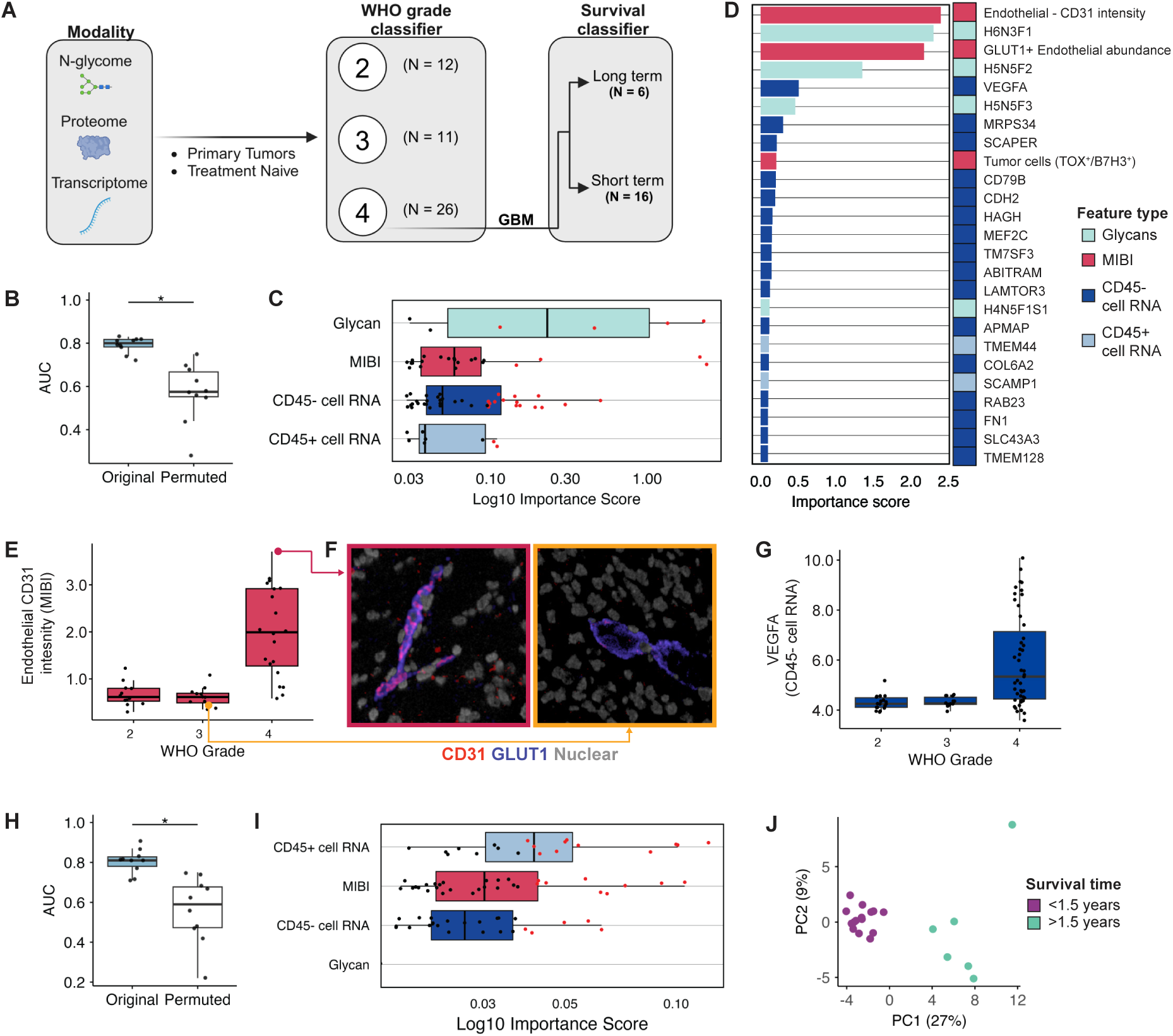
Multi-omic classification of glioma grades and survival features. **(A)** Schematic of the study design for classifying glioma patients by WHO grade (2, 3, 4) and distinguishing between short-term and long-term survivors in GBM using multi-omic data (transcriptome, proteome, N-glycome) **(B)** Boxplot showing the Area Under the Curve (AUC) for the WHO grade classifier with original and permuted labels. **(C)** Top 75 Log10 importance scores for different feature types (glycans, CD45+ and CD45-cell RNA, and MIBI) used in the WHO grade classifier. **(D)** Top 25 feature importance ranking. Glycan (Cyan), MIBI (red), CD45-cell RNA (dark blue) and CD45+ cell RNA (light blue) **(E)** Boxplot of endothelial cell CD31 intensity across WHO grades, with high and low representative image of CD31 expression. Whole image shows CPM with endothelial cells colored in red **(F)** Inset shows raw CD31 marker expression **(G)** Boxplot showing VEGFA RNA expression levels across WHO grades for CD45-cell populations. **(H)** Boxplot showing AUC values for the survival classifier with original and permuted labels **(I)** Top 75 Log10 importance scores for different feature types (glycans, CD45+ and CD45-cell RNA, and MIBI) used in the WHO grade classifier **(J)** Principal Component Analysis (PCA) plot using top 75 features from multi omic classifier. Each dot represents a patient. blue for long term survivors (>1.5 years) and red for short term survivors (<1.5 years).

While tumor grading across diverse cancers, including gliomas, breast, and lung tumors, is typically determined by morphological changes, mutational status, and vascular proliferation, the molecular determinants of patient survival often diverge from these grading criteria^75^. To that end, using a similar approach, we trained a separate model for classifying short-term vs long-term survival in treatment-naive primary GBM to understand what aspects of the glioma TME separate these groups (**Figure 7A**). This model exhibited similar performance to the one above with an AUC of 0.81 and a permuted baseline of 0.59 (**Figure 7H**). However, the top 75 features for each model markedly differed with only the ratio of B7H3+ to TOX+ tumor cells being selected in both (**Tables S4 and S5**). In contrast to the grade-specific model, none of the top 75 survival-associated features included glycans (**Figure 7I**). Similarly, while transcripts from immune-rich regions had the lowest median importance for grade they had the highest median importance for survival predictions. This suggests that within the most aggressive disease, GBM, tumor immune responses are the predominant driving factor of survival. Transcripts such as MCM2, KIF3A, THOC6, and TMED3, all known to play a major role in tumor progression were among the most important features from immune-rich regions (**Figure S7B, Table S5**). To further validate this association by restricting the search space further, we confirmed that the top 25 features (**Figure S7B**) from the classifier were truly distinguishing between short-and long-term survivors, we performed PCA. The results showed a clear separation of the two groups along the PC1 axis, which accounted for 27% of the variance (**Figure 7J**). Together, these results demonstrate that while tumor glycan and vascularization features are most relevant for distinguishing / classifying tumor grade, immune-related transcriptional programs play a more critical role in driving survival signals, underscoring the value of multi-omic profiling in categorizing disease state progression and understanding and possibly perturbing patient outcomes.

## Discussion

Gliomas are among the most aggressive tumors with the lowest overall survival (OS) rates of all cancers^1^. While significant improvements in OS have been achieved with immune or tumor targeted-therapies in extracranial malignancies like melanoma or lung adenocarcinoma, comparable progress in gliomas remains elusive^33,76,77^. This lack of progress suggests that mechanisms of tumor persistence in gliomas differ fundamentally from those occurring outside the blood brain barrier. To gain insights into why these approaches have stalled and how future therapies could circumvent these issues, we assembled a multicenter glioma cohort from 310 patients, representing 677 samples, to provide a unified, comprehensive view of this unique TME **(Figure 1)**. Multiomic integration of spatial proteomics, transcriptomics, and glycomics revealed systematic differences in the structure, composition, and phenotype of gliomas with respect to tumor type and grade, a rationale for why targeted therapies have failed to improve OS, and new insights into how the tumor glycocalyx could be driving severity in high grade lesions. To enable future work to build upon these findings, we have made the data, analysis, and clinical endpoints freely available, without restriction, via an interactive web portal at www.bruce.parkerici.org **(Figure 1)** where users can browse annotated single cell data and view spatial proteomics images. This resource is also hosted on Bioimage Archive and GitHub where it can be downloaded in its entirety.

In previous work, we found that TME organization and morphology can be used to construct accurate trajectories of immune recruitment, infer immune phenotype, and to predict OS in both invasive and pre-invasive breast cancer^65,78^. Likewise, work by others have found this concept of spatial rulesets generalizes to other malignancies like melanoma and head and neck cancer as well^61,79^. In contrast, gliomas are highly diffuse and unstructured, often lacking the type of consistent spatial organization that is well suited for automated, extensible, and accurate scoring needed for developing these spatial rulesets using conventional tools.

To address these challenges on a larger scale with the less-structured TME of glioma, we used QUICHE to uncover subtle structural trends that were differentially expressed between patient groups^63^. For example, comparing IDH-mutant LGG structure in patient-matched primary and recurrent lesions revealed consistent and significant changes in how tumor and immune cell subsets localized with one another **(Figure 5)**. We would like to highlight two points from this analysis. First, while the TME structure differed significantly in primary and recurrent lesions, the prevalence of tumor and immune cell subsets did not. This means that the changes in cell localization observed here are likely due to corresponding differences in TME signaling rather than shifts in cell prevalence that could drive more happenstance interactions. Second, cells located within these differential niches were phenotypically distinct from cells outside of them. For instance, TIM3 and IDO1 were elevated in niches differentially enriched in primary tumors but not in recurrences. Intriguingly, previous work found the spatial organization of cells expressing TIM3 and IDO1 to be predictive of outcome in ovarian cancer, suggesting that the spatial organization of these cells are a surrogate for shifts between productive and tolerogenic immune responses. In contrast, when comparing on-therapy samples from patients receiving IO with those who received SOC, no structural differences were found. These findings suggest that while IO therapy may alter the functional status of certain immune populations, the lack of change in spatial organization could indicate these effects are transient and insufficient for recruiting durable antitumor immunity.

Tumor-targeting therapies such as small molecule EGFR inhibitors, monoclonal antibodies targeting B7H3, or CAR-T therapies against B7H3 and EGFR are another class of interventions that are being explored^10,50,80^. Like immunotherapy, these approaches have been effective in some tumors outside the brain but thus far in early clinical trials have not improved OS in glioma^4^. To understand why this might be, we conducted the first quantitative analysis of TA for seven targets currently in clinical trials **(Figure 2 and 3)**. This analysis revealed three insights for understanding why this approach has not yet been successful and how drug regimens might be optimized in future studies. First, ∼57% of gliomas had a high proportion of tumor cells (>30%) that were pan-negative for all TAs quantified here, suggesting that these patients may not benefit from these therapies.

Second, for lesions with higher proportions of TA-positive cells, B7H3 and EGFR were the most frequently expressed across all gliomas. Tumor cells in ∼69% of GBM patients expressed one of these antigens in comparable proportions to CD19 in B-cell malignancies (>72% antigen positive tumor cells). However, B7H3 and EGFR also exhibited the highest degree of cell-to-cell variation in expression (6-12 fold) **(Figures 4)**. Given that low antigen copy number has been shown previously to be associated with therapeutic failure in solid tumors, the proportion of glioma cells with weak antigen expression could be a determinant here as well^81^. Lastly, personalized therapy targeting one or more antigens could provide a means of improving tumor targeting and mitigating antigen escape^57^. For example, early phase clinical trials using bispecific CD19/20 CAR-T to treat Non-Hodgkin Lymphoma have shown >2 times the PFS compared to using CD19 CAR-T alone^82^. Based on our results, B7H3 and EGFR should offer the most coverage and overlap in GBM, Astrocytoma and oligodendroglioma. In contrast, adult PXA patients showed high tumor cell coverage with just B7H3, potentially making them ideal candidates for monotherapy. Still, given the broad diversity of antigen expression (**Figure 2–4**) and the significant influence of immune microenvironment on patient survival (**Figure 7**), effective long-term GBM treatment will likely depend on strategies that harness and activate endogenous adaptive immune responses.

Recent advancements in glioma molecular classification, underscore the need to explore novel biological dimensions beyond the proteome and transcriptome to fully capture tumor heterogeneity and progression^83^. To understand how glioma tumor *N*-glycans are involved in driving disease progression, we used a selective PNGaseF catalyzed cleavage assay with MALDI-IMS for label-free, *de novo* detection of these structurally diverse posttranslational modifications^68^. *N*-glycans are present on most membrane proteins where they comprise a large portion of glycocalyx, a viscous carbohydrate coating surrounding the cell membrane that is involved in regulating a broad range of cell-cell interactions^84^. Aberrant tumor glycosylation is a cancer hallmark that was first identified over 50 years ago^85^. Since that time, innovations in chemical biology and mass spectrometry have revealed the startlingly central role it plays in nearly every facet of disease progression^86^. For example, TME sialylated glycans drive immune evasion by binding Siglecs present on both lymphoid and myeloid cells^34–36^.

Importantly, healthy brain tissue has distinct N-glycosylation patterns critical for normal neuronal function and immune regulation^87^. Various sialylated structures in different brain subregions modulate neuronal activity and intercellular communication, reflecting the tight regulation of the brain’s structural and immune milieu^88^. Disruption of these glycan profiles can have far-reaching consequences, underscoring how aberrant sialylation in gliomas may potentiate immune evasion and malignant infiltration.

Recognizing this, we quantified N-glycan class enrichments across glioma grades and found that sialylated glycans were significantly enriched in grade 4 gliomas, where immune cells represent a larger proportion of the tumor mass (**Figure 6**). To understand how these changes in the tumor glycocalyx impact glioma composition and function, we correlated these findings with co-registered spatial proteomics and transcriptomics. Tumor sialylation correlated with preferential increases in specific immune subsets, including CD209+ DC/Mac, CD4+ T cells, and Tregs (**Figure S6**). In line with these findings, previous studies have demonstrated that DC uptake of sialylated antigens induces tolerogenic states that preferentially polarize naïve CD4+ T cells toward Tregs^89^.

Expanding on this, we trained a classifier on all three spatial modalities to simultaneously determine contrasting glioma features that were predictive of grade and clinical outcome (**Figure 7).** Tumor glycans and features relating to neovascularization (i.e. CD31 endothelial expression, VEGFA transcript abundance) accounted for the top five most predictive of glioma grade. In contrast, these same spatial glycomic signals provided almost no value for predicting OS in patients with GBM. Conversely, while spatial transcriptomics of immune-enriched CD45-positive regions were the least valuable for predicting grade, they were the most valuable for predicting GBM outcomes. These results indicate the divergent properties, specific to high grade disease, should not be assumed to drive clinical outcome as well.

Taken together, this study provides a comprehensive view of the multi-omic landscape of human gliomas. We established baseline measurements of tumor antigen variability, immune cell landscapes, and the effects of therapeutic perturbations on the TME. Additionally, for the first time we uncovered dynamic relationships between glycosylation, cellular features, and transcriptomes in humans, creating a unified dataset that serves as a valuable resource for understanding glioma biology and advancing translational research for this disease.

Going forward, multiple directions have emerged that future studies can build upon to deepen our understanding of glioma pathophysiology. First, potential off-target consequences of B7H3-directed treatments warrant scrutiny, given its prominent expression on non-tumor cells within the brain. Additionally, strategies to enhance treatment efficacy should account for the notable variability of tumor antigens at the single-cell level. Longitudinal studies will help us understand how the tumor glycocalyx and TME architecture change over time, which could explain why current therapies have limited effects. Using these insights in new clinical trials, especially those testing bispecific or multiantigen-targeted treatments that account for individual glycocalyx and antigen profiles, may help overcome the barriers that have held back glioma immunotherapy and targeted therapies.

### Limitations of Study

Despite the broad and diverse cohort in this study, having a larger sample size for each clinical factor would have further increased the confidence of our findings. While we included longitudinal sampling for some IDH mutant LGG lesions, having patient-matched primary and recurrent high-grade samples would have provided even deeper insights into tumor evolution. The nature of this study was entirely retrospective, and thus our findings, however significant, are primarily correlative. Our use of TMA, though guided by expert pathologists to capture areas rich in tumor and immune cells, inevitably limited the tissue coverage per patient and may have missed important regions. In terms of single-cell analysis, although we measured a relatively large panel of markers, future work could refine this panel to include additional markers for cytokines for better immune cell functional characterization, as well as more robust markers for tumor cells. Our spatial whole transcriptome profiling offered a perspective on immune-rich and immune-poor regions, but a single-cell RNA approach might have yielded complementary data with more cellular granularity more suited for multi-omic integration with our single-cell protein data. Finally, practical constraints such as multi-site coordination and patient confidentiality meant we could not obtain a complete clinical history for individuals, indicating that, while our dataset is comprehensive, there is still room for improvement in both coverage and annotation.

## Methods

### Study design and sample collection

This cohort was assembled by combining tissue samples from several clinical trials and institutional archives spanning multiple disease types, ages, treatment conditions, and WHO grades. Multiple regions were collected per patient based on location and CD45/CD3 IHC for immune rich regions. All samples were chosen under the supervision of a pathologist. A total of 677 cores from 310 unique patients were included on the 16 tissue blocks.

Phase I trial for GBM Response to Pembrolizumab (NCT02852655) of surgically accessible recurrent/progressive glioblastoma that received 200 mg pembrolizumab (MK3475) intravenous infusions. All patients provided written informed consent; the study was approved by institutional review boards at all sites (Dana-Farber Cancer Institute; Huntsman Cancer Institute; M.D. Anderson Cancer Center; Massachusetts General Hospital; Memorial Sloan Kettering Cancer Center; University of California, Los Angeles; University of California, San Francisco) and conducted according to the Declaration of Helsinki.

The LGG vaccine trial was made up of two phase I trials (NCT02924038, NCT02549833) (n = 53). Trial 1 was a pilot, randomized, two arm neoadjuvant vaccine study in human leukocyte antigen-A2 positive (HLA-A2+) adults with World Health Organization (WHO) grade II glioma, for which surgical resection of the tumor is clinically indicated. 4.96mg IMA950 and 1.4mg poly-ICLC administered as one formulation followed by/or without 3mg/kg Varlilumab infusion (intravenously) ∼3 weeks before the date of scheduled standard-of-care surgery. Patients continue receiving IMA950/poly-ICLC subcutaneous injections every week leading up to surgery. Trial 2 was a pilot neoadjuvant vaccine study in adults with WHO grade II glioma, for which surgical resection of the tumor is clinically indicated. GBM6-AD lysate protein and poly-ICLC administered as one formulation every week leading up to standard-of-care surgery to grade II glioma ∼3 weeks or was given post-surgery. Both studies were approved by the IRB at UCSF and were conducted according to the Declaration of Helsinki, all patients provided written informed consent.

The trial combining IL13Rα2-CAR-T cells with checkpoint inhibition (NCT04003649) is a phase 1, single-center, randomized study, comparing CAR T cells plus nivolumab with (Arm 1) and without (Arm 2) neoadjuvant treatment with nivolumab and ipilimumab seven days before surgery. The primary objectives are to establish the safety and feasibility of administering 50 x10^6^ IL13(EQ)BBζ/CD19t+ T cells (100 x 106 T cells total) for 4 week cycles (n = 6 research subjects per Arm) via dual delivery (administered through both the ICT and ICV catheters) in combination with IV administration of checkpoint inhibitors in participants with recurrent glioblastoma and IDH-mutant astrocytoma (grade 4). Samples collected were prior to CAR-T administration.

Treatment naive GBM samples tissues were acquired from Stanford Health Care’s tissue repository (n= 86) in which of the tumor samples there are, grade 2 astrocytoma (n = 5), grade 2 oligodendroglioma (n =5), grade 3 astrocytoma (n = 9), grade 3 oligodendroglioma (n =3), grade 4 astrocytoma (n = 8), and grade 4 GBM (n= 32). A select set of patient samples were labeled as either core (n = 21), core-to-infiltrating (n = 8), or infiltrating edge (n = 14) depending on the location of the sample within the tumor.

Pediatric H27M+ diffuse midline gliomas (DMG) and other pediatric high-grade gliomas (pHGG) were acquired from Children’s Hospital of Philadelphia (CHOP) (n =156). Of these, TMAs included tumors designated as pediatric astrocytoma (n = 24), pediatric midline glioma (n = 25), pediatric GBM (n = 33), pediatric ganglioglioma (n = 2), pediatric glioma (n = 16), pediatric high grade gliomas (n = 8), pediatric PXA (n = 8), and pediatric thalamic glioma (n = 2), as well as breast carcinoma (n = 4) and colon carcinoma (n = 4) as control tissues. More specific diagnostic categorization of the pediatric cases, including integration of molecular diagnostic information when available, can be obtained on request from the Children’s Brain Tumor Network (CBTN, https://cbtn.org/).

### Control TMA construction

We constructed a control tissue microarray (TMA) to assess slide-to-slide staining variability. The TMA comprised five 1.5mm control tissue cores sourced from archival FFPE specimens provided by the Stanford pathology department, including duplicate cores of tonsil, lymph node, and placenta, plus an extra tonsil core for asymmetry. Serial recuts of the control TMA, as well as the cohort TMAs, were placed on the same slides. Additionally, a larger control TMA was constructed of 2 mm cores of normal brain, tonsil, spleen, placenta, high grade glioma, thymus, lymph node, testis, kikuchi disease, colon carcinoma, cholangiocarcinoma, liver, low grade glioma, melanoma, ependymoma, breast carcinoma, ovarian cancer, lung scc, DIPG.

### MIBI staining

#### Panel construction

Most antibodies used in this study were previously validated for MIBI-TOF^65,90,91^. Newly selected target antibodies were validated via immunohistochemistry to confirm appropriate staining in control tissues. All antibodies were metal-labeled using the Ionpath conjugation kit (IonPath, Menlo Park, USA) according to the manufacturer’s protocol. Calprotectin and Mast Cell markers (Chymase and Tryptase) were conjugated to Ga69 and Ga71 respectively using monomeric maleimido-mono-amide NOTA. To prevent degradation and prolong shelf life, labeled antibodies were lyophilized individually with 100 mM trehalose in 1 µg or 5 µg aliquots. Titer optimization for new targets was performed using serial dilutions (1 µg/mL, 0.5 µg/mL, 0.25 µg/mL, and 0.125 µg/mL) or the recommended titer for previously validated MIBI-TOF antibodies.

#### Cohort staining

To minimize batch effects, all TMAs were stained using two mastermixes: one for the tumor panel and one for the immune panel. Serial TMA slices were stained separately for each panel. Fresh antibody aliquots were reconstituted and combined into the mastermixes. To reduce errors, staining was performed in pairs, with one person reading and the other pipetting. Detailed procedures are available in our methods publication, with a step-by-step guide outlined below^28^.

#### Interactive protocols

Reagent preparation: https://www.protocols.io/view/mibi-and-ihc-solutions-261geo7wyl47/v1 IHC staining: https://www.protocols.io/view/ihc-staining-x54v9moxmg3e/v1 MIBI staining: https://www.protocols.io/view/mibi-staining-dm6gprk2dvzp/v5

Sequenza staining: https://www.protocols.io/view/staining-sequenza-6qpvrdeo2gmk/v1 Antibody lyophilization: https://www.protocols.io/view/antibody-lyophilization-kxygxex5kv8j/v1MIBIdatageneration

#### MIBI run setup

MIBI data was acquired using the MIBIScope instrument (IonPath, Menlo Park, USA). TMA cores were labeled by row and column for easy mapping to metadata. Each core was acquired on two slides, one for the tumor panel and one for the immune panel, with screen captures taken to align regions between slides. Automated checks, with user confirmation, corrected naming errors in row and column assignments. Core order was randomized before acquisition to minimize batch effects from instrument drift^92^.

#### Acquisition settings

Consistent settings were applied across all MIBI data acquired in this study. Field of view (FOV) size was selected based on sample availability. When feasible, FOVs measuring (800 µm)^2^ and (2048 pixels)^2^ were captured. In cases of limited tissue, an FOV size of (400 µm)^2^ and (1024 pixels)^2^ was used instead. A custom preset with a beam current of 8.5nA and dwell time of 0.58ms was applied to optimize acquisition speed and image clarity. All default background correction and noise removal settings were disabled.

#### Image compensation with Rosetta

We used our previously described Rosetta Matrix Compensation as a method for correcting contamination and background across channels in MIBI data^64^. In summary we empirically determined signal overlap for each source-output channel pair in order to create a matrix of all channel overlaps and use this as a template to subtract known overlapping source markers from their target counterpart, similar to flow-cytometry compensation.

#### Intensity normalization using median pulse height

We used our previously described Intensity normalization using median pulse height (MPH) as a method for correcting for decreased sensitivity across a run in MIBI data^64^. In summary we created an exponential curve with a PMMA (polymethyl methacrylate) sample, linking MPH to a normalization coefficient. Each channel then has a polynomial function that is fit individually per mass over the run that ignores outliers in order to produce a predicted MPH for that image. This MPH is then mapped to the earlier described exponential curve to get a normalization coefficient for each image. We then use this coefficient to normalize the data, correcting for drift and restoring consistent signal levels throughout the run. This approach mitigated the observed decrease in signal over time, preserving data accuracy.

### MIBI data QC

To detect potential batch effects, we assessed slide-to-slide variation. For batch effects across slides, control samples on each slide served as references, allowing us to calculate changes in the mean of non-zero pixels for each channel in each control sample.

### Cell segmentation

We segmented all images in the cohort using the previously described Mesmer model^93^. Mesmer is a pre-trained deep learning model that utilizes two input channels: a nuclear marker and a membrane marker. For the nuclear channel, we used the HH3/dsDNA channel, and for the membrane channel, we combined the CD14, CD45, CD8, and HLA1 channels.

### Cell clustering

#### Pipeline overview

Pixie is an unsupervised pixel clustering algorithm that uses the subcellular localization of proteins to improve cell classification from multiplexed image data^94^. We used pixie to phenotype cells in a four-step process. First, pixels were over-clustered in each image by training a self-organizing map (10 × 20 grid or 200 clusters) on the phenotypic lineage marker pixels across all images for both panels^95^. For the immune panel we used Calprotectin, CD3, Chym/Tryp, CD68, CD4, CD8, CD208, CD141, CD209, CD11b, CD14, CD123, CD20, CD206, CD31, CD45, CD133, CD163, GFAP, GLUT1, FoxP3, HLADR, NeuN, Olig2, TMEM119. For the tumor panel we used CD4, FOXP3, CD31, NeuN, CD8, CD3, CD163, TMEM119, HLADR, CD14, CD45, ApoE, NG2, B7H3, VISTA, EGFR, HER2, EGFRvIII, EphA2, CD70, GPC2, H3K27M, GM2 GD2, CD133, GFAP, HLA1, Olig2, IDH1 R132H, CD47. These pixel clusters were then grouped into 20 pixel metaclusters using consensus hierarchical clustering^96^. Next, for each cell in the image, we counted the proportion of pixels in each pixel metacluster to generate a *cell* × *pixel metacluster* matrix. Following the same procedure, cells were then over-clustered by training a self-organizing map (10 × 20 grid or 200 clusters) on this pixel metacluster matrix, then grouped into 20 cell metaclusters using consensus hierarchical clustering. We validated these predictions through manual inspection of cell type-to-marker expression using Adobe Photoshop. Pixie was implemented using the ark-analysis v0.7.0 package in Python.

#### Cluster inspection and cleanup

To confirm accurate clustering, tiles of all images within each panel were constructed. Cell assignments were manually verified by looking at images through Adobe Photoshop. Assignments were viewed and confirmed by toggling on the appropriate image channels, which were then overlaid with the segmentation boundary and cluster assignments. Systematic errors in clustering were then addressed by reclustering based on marker specific thresholds.

#### Combining Immune and Tumor Panels

To integrate immune and tumor panels collected from separate serial slices, each with its own cell table, a comprehensive table encompassing all tumor and immune markers was constructed.

For each field of view (FOV), the immune FOV and tumor FOV were spatially aligned to ensure accurate integration.

Each FOV was evaluated and categorized into one of three groups: (1) already aligned, (2) requiring manual alignment, or (3) unable to align. For FOVs requiring manual alignment, cells from both panels were plotted, and one FOV was shifted along the x and y axes to match the spatial layout of the other. Post-alignment, cells lying outside the aligned space were excluded. FOVs that could not be aligned, due to discrepancies in serial slices, variations in FOV sizes, or missing FOVs, were excluded from further analysis.

From the immune panel (288,224 cells), immune cells, endothelial cells, and neurons were retained, while tumor cells were excluded. From the tumor panel (869,069 cells), only tumor cells were retained, with immune cells, endothelial cells, and neurons removed. This resulted in a single final combined panel (1,157,293 cells) prevented double counting and ensured the accurate integration of immune and tumor cells.

### Feature extraction pipeline

#### Pipeline overview

Feature extraction pipeline incorporates a diverse array of features to thoroughly characterize the tumor-immune microenvironment. For each sample in our cohort, we computed all features, such as cell composition, functional marker expression, cell density, cell ratios, transcriptome gene counts, and glycome relative intensities. Each feature was calculated at the image level. To ensure accuracy, we applied minimum cell count thresholds for many of these calculations, omitting any features that did not meet these thresholds in a given image. Once generated, all features were transformed into a standardized format and consolidated into a single data frame for downstream analysis. Details available at https://github.com/angelolab/publications/tree/main/2024-Piyadasa_Oberlton_etal_Glioma.

#### Aggregating computed features

Once the image features were calculated, we integrated them into a unified data structure for downstream analysis. Each feature was assigned a descriptive name, along with metadata specifying the image compartment where it was calculated, the cell types and/or markers involved, and its overarching feature category.

### Gene Set Enrichment Analysis (GSEA)

For the proportion of either GM2/GD2 or B7H3 out of all tumor cells, the median value was calculated across all samples. Log2 fold changes were computed for each sample by dividing the feature value by the median value, followed by sorting in descending order to generate a ranked list of samples. Samples were then grouped into two categories based on tumor region: “Tumor Core” and “Tumor Infiltrating.” Each group was treated as a gene set for GSEA. The ranked list of samples, based on log2FC values, was used as input for GSEA. The minimum and maximum sizes of gene sets were automatically calculated based on the group sizes. Enrichment scores were then calculated for both tumor regions using R package fgsea v1.30.0.

### QUICHE Analysis

We performed Quantitative InterCellular Niche Enrichment (QUICHE) analysis to identify differentially enriched niche neighborhoods across conditions (e.g. LGG primary vs recurrent, tumor core vs tumor infiltrating edge, and GBM vs Grade 4 astrocytoma)^63^. In summary, for each condition comparison, we identified cellular niches according to spatial proximity within each image, performed distribution-focused downsampling to select a subset of niches per patient sample, constructed a k-nearest neighbor (k-NN) graph for differential neighborhood enrichment testing, and labeled niche neighborhoods according to the most abundant cell types (knn = 50, radius = 200, k_sim = 100, looking at top 3 cell types per niche, downsample was determined to be within 1 standard deviation below the mean, and adjusted depending on the spread of cells per FOV). Niche neighborhoods were considered significant if they had a spatial false discovery rate (FDR) < 0.10. QUICHE was implemented using the Python package at https://github.com/angelolab/publications/tree/main/2024-Ranek_etal_QUICHE.

### QUICHE vs Bulk Functional Marker Analysis

To compare the functional marker abundance within QUICHE neighborhoods versus on the bulk scale we started by first calculating, for each functional maker, the percent positive cells in each cell type within a differentially expressed neighborhood. From there we then aggregated all cells of the same type across all differentially expressed niches and calculated the mean percent positivity for each functional marker. We separately calculated, for each functional marker, the bulk cell percent positivity for all cells in each condition of the comparison criteria. For each condition we calculated the Log2FC between the QUICHE percent positivity and the bulk percent positivity. If <0.05 of bulk cells were positive for a given functional marker the Log2FC was filtered out. In order to avoid division by zero a pseudo count of 1e-6 was added to all values. In order to keep from large Log2FC values a cap was created at-5 and 5.

### Random Forest Model

#### Data Import and Preprocessing

We filter the data based on criteria such as modality, immunotherapy status, disease type, patient age, and tumor region to ensure that only relevant variables are included in the analysis. Further data cleaning is performed by removing highly correlated features. A correlation matrix is calculated to identify highly correlated features. Features exceeding a specified correlation threshold (0.6 for who grade and 0.72 for survival status in order to maintain a similar number of features between the conditions) had the feature with the largest mean absolute correlation removed, retaining only those that contribute unique information to the analysis., eliminating redundancy in the dataset. We also exclude features with a large proportion of >50% missing values and features in which all values are the same.

#### Feature Harmonization and Normalization

Each feature is normalized (Z-scored) to facilitate comparisons. For datasets with multiple fields of view (FOVs), feature values are averaged across FOVs to ensure uniform representation.

#### Model Setup

Using the processed data, a random forest classifier is trained to predict survival status (i.e. short vs long) or who grade (i.e. 2, 3, 4). We implement k-fold cross-validation (k = 3) to evaluate model performance. To evaluate performance, we compared the predictions to the ground truth using the area under the receiver operator curve (AUC). We performed 3-fold cross validation in each of 10 runs to obtain a distribution of predictions.

#### Model Evaluation and Feature Importance

For each training iteration, AUC values are recorded to assess model performance. The median importance of each feature across multiple runs is calculated using the Gini importance metric, and the top features are identified based on this metric.

#### Randomized Control

To validate model performance, labels are randomly shuffled outcome labels without replacement and the model is re-run to compare AUC values against the original (non-randomized) model, random forest is implemented in R using caret v.6.0.94. A t-test is implemented in R using stats v.4.4.1 to determine statistical significance between the original and randomized AUC values, providing a measure of model reliability.

#### Principal Component Analysis (PCA)

Principal components analysis (PCA) was performed on the top 20 features for visualization in R using stats v4.4.1.

### RNA-Glycan co-occurrence analysis

In order to investigate the correlation between the gene expression of enzymes involved in glycan generation and the abundance of glycans across different GBM stages, we conducted an analysis on differentially expressed genes and glycans.

Initially, we normalized the expression data for these genes and glycans by Z-scoring each gene or glycan’s expression across GBM stages. Thereafter, k-means clustering was applied using k=6, a value driven by minimal k to account for all potential trends across the three stages of GBM. This ensured sufficient classes to account for any trends observed within our dataset, while minimizing clustering complexity.

Using literature references, glycans and genes were each assigned a glycan type, such as bi-antennary or fucosylated. It’s noteworthy that a gene or glycan could be assigned to multiple types. Following this assignment, we quantified, for each glycan type, instances where both a glycan and an enzyme of that type shared the same k cluster.

For establishing a comparative basis, we generated a null distribution by randomly assigning genes and glycans to types 1,000 times. The previously calculated quantity was then Z-scored against this null distribution. A high Z-score indicates a glycan type with a more correlated expression of genes and glycans than one would expect by chance. This method accounts for the overall expression trends observed in our dataset as given by the distribution between k-clusters.

### N-glycan MALDI imaging mass spectrometry of FFPE tissue slide preparation

Tissue TMA slides were prepared for N-glycan imaging mass spectrometry analysis using a standardized tissue preparation workflow which has been previously published for FFPE tissues^97^. Briefly, tissue slides were dewaxed with xylene and rehydrated using a gradient of ethanol in a linear stainer (Leica Biosciences) programmed to 3 dips per wash for 30 seconds each. Antigen retrieval was performed in citraconic anhydride buffer at pH 3 for 30 min in a heating chamber at 95 °C. Slides were briefly dried under vacuum and subject to enzyme treatment using HTX M3+ Sprayer (HTX Technologies, Chapel Hill, NC).5 passes of PNGaseF PRIME enzyme (N-Zyme Scientifics, Doylestown, PA) at 0.1 µg/µL was applied at a rate 25 µL/min with a velocity of 1200 mm/min and a 3 mm offset at 10 psi and 45 °C. Slides were incubated in prewarmed humidity chambers for 2 h at 37 °C. After PNGaseF digestion, 7 mg/mL CHCA matrix in 50% ACN/0.1% TFA was applied to the slides at a rate of 100 µL/min with a velocity of 1300 mm/min and a 2.5 mm offset at 10 psi and 79 °C using the same sprayer. Slides were stored under vacuum until analysis^97^.

### MALDI-MSI Data Acquisition and Processing

All samples were analyzed with a timsTOF fleX MALDI-2 mass spectrometer (Bruker Corporation, Billerica, MA). The following parameters were used: mass range 800-4000 m/z, positive ion mode, 10 μm field size and 200 shots per pixel. Raw imaging data was processed in SCiLS Lab version 2024b Pro (Bruker Corporation, Billerica, MA). In SCiLs Lab, N-glycan spectra were normalized by total ion count and converted to vendor-neutral imzML format. The imzML files were parsed using pyimzML in Python, and the expression at each m/z peak was extracted as single-channel TIF images.

### Nanostring DSP Sample Preparation

Formalin-fixed, paraffin-embedded (FFPE) samples were prepared manually according to NanoString GeoMx RNA-NGS protocols. Tissue microarrays (TMAs) were arranged to fit the largest dimensions allowable for imaging on the DSP system. Five-micrometer sections were placed on positively charged slides, baked, deparaffinized, and then sequentially washed in ethanol and PBS. Antigen retrieval was performed with Tris-EDTA (pH 9.0) in a pressure cooker for 10 minutes at 100°C, followed by a PBS rinse. The samples were then incubated with Proteinase K (1 µg/mL) for 15 minutes at 37°C and washed again in PBS.

Afterward, the slides were hybridized overnight at 37°C with human WTA at the recommended concentration. HybriSlip Hybridization Covers (Grace BioLabs) were applied during this step. Coverslips were detached by soaking the slides in 2× SSC plus 0.1% Tween-20, and any unbound probes were removed by two 25-minute washes in 50% formamide/2× SSC at 37°C, followed by a brief wash in 2× SSC.

For staining with morphology markers, slides were first incubated in blocking buffer at room temperature for 30 minutes in a humid chamber. They were then exposed to 500 nM SYTO13 and fluorescently labeled antibodies (CD45, GFAP) for 1–2 hours. A final wash in 2× SSC was performed before loading the slides onto the GeoMx DSP instrument.

### Nanostring DSP acquisition

DSP experiments were conducted in alignment with the NanoString GeoMx-NGS DSP Instrument manual and established protocols^98^. In brief, slides were imaged across four fluorescence channels (FITC/525 nm, Cy3/568 nm, Texas Red/615 nm, and Cy5/666 nm) to visualize the selected morphology markers. Regions of interest (AOIs) were defined and collected for all samples, with segmentation driven by CD45 high and CD45 low expression using the DSP auto-segmentation tool and manually adjusted parameters. The selected AOIs were illuminated, and the released tags were deposited into 96-well plates following the established protocol.

### Sequencing and data cleanup

Following the NanoString GeoMx-NGS Readout Library Prep guidelines, the dried DSP aspirate was reconstituted in 10 µL of DEPC-treated water, and 4 µL of this solution was used for PCR with NanoString SeqCode primers. These primers both amplify the tags and incorporate Illumina adaptor sequences along with sample-specific barcodes. PCR products were pooled in fixed volumes and then purified twice using AMPure XP beads (Beckman Coulter). The final libraries were sequenced exclusively on an Illumina NovaSeq 6000 using 27 × 27 paired end reads.

FASTQ files were handled with the NanoString GeoMx NGS Pipeline v2.0 or v2.2. Low-quality bases and adaptor sequences were trimmed, and paired-end reads were stitched together. Barcode and UMI regions were identified, allowing up to one mismatch for barcode matching. Reads sharing the same barcode were deduplicated by UMI. The raw read count correlated strongly with unique UMIs across AOIs (Supplemental Figure S9), supporting a generally consistent library amplification. Nevertheless, all analyses presented here employed UMI-deduplicated counts to account for potential PCR bias.

Count data were processed and normalized using GeoMxTools R package v1.0 (https://bioconductor.org/packages/release/bioc/html/GeomxTools.html). Specific parameters and detailed used can be found in https://github.com/angelolab/publications/blob/main/2024-Piyadasa_Oberlton_etal_Glioma/Data_Analysis/2_NS_DSP_cleanup.R

## Resource availability

### Lead contact

Requests for further information and resources should be directed to and will be fulfilled by the lead contact Michael Angel.

### Materials availability

All unique/stable reagents generated in this study are available from the lead contact with a completed materials transfer agreement.

### Data and code availability

To provide broad access to our multi-omic data set (including single cell MIBI image overlays) and clinically annotated metadata, we have created an interactive online portal, www.bruce.parkerici.org. This portal further facilitates direct download of all raw data tables and features calculated. Raw MIBI images and segmentation files are deposited at BioImage Archive (https://www.ebi.ac.uk/biostudies/bioimages/studies/S-BIAD1579) for easy access.

Code to generate all figures is available at https://github.com/angelolab/publications/tree/main/2024-Piyadasa_Oberlton_etal_Glioma. Low-level processing code can be found at: https://github.com/angelolab/toffy. Spatial analysis code can be found here: https://github.com/angelolab/ark

## Acknowledgements and funding

We thank the patients and their families for participating in all the trials. We thank Parker Institute for Cancer Immunotherapy (PICI) that supported the multi-site collaboration, funding the project and the development of the web resource. We also thank Alliance for Cancer Gene Therapy for their funding support for additional samples, as well as their scientific oversight. We acknowledge resources provided by the UCSF Brain Tumor SPORE Biorepository and Pathology Core. Dr. Piyadasa was supported by the Canadian Institutes of Health Research (CIHR) postdoctoral fellowship (FRN: 176490). N.F.G. was supported by NCI CA246880, NCI CA264307, and the Stanford Graduate Fellowship. Dr. Phillips was partially supported by the National Institutes of Health (R01 NS131474). NIH/NCI P50CA097257 for JJP. NIH/NCI U01 CA242096 for RRD.

Dr. Oldridge was supported by NHLBI R38 HL143613, NCI T32 CA009140, and a bridge fellowship from the Parker Institute for Cancer Immunotherapy and V Foundation for Cancer Research.

## Author contributions

H.P., B.O., S.B. and M.A. conceived the project.

R.P., H.O., J.P., D.O., C.B., M.B, K.C., F.G., P.P., F.F., A.T., S.F. and M.G.H collected and processed FFPE tissues, clinical metadata and provided expert guidance and feedback throughout the project.

E.Y., M.T., M.S., S.B. and S.L. facilitated the multi institutional collaboration and provided funding for the project.

H.P., E.Y., M.T., C.S. and S.V. constructed the web portal and deposited data.

H.P., B.O., I.A. C.C.F., M.Bo., R.K., and M.R. generated and processed MIBI, MALDI and NanoString DSP data.

H.P., B.O., J.R., I.A., K.L., M.A., C. L., N.F.G., E.M., A.K., C.S. and M.S. analyzed multi omic data.

H.P., and B.O. generated figures.

H.P., B.O., S.B. and M.A. wrote the manuscript.

S.B. and M.A. supervised the project.

R.D. provided expert feedback on MALDI analysis. All authors provided feedback on the manuscript.

## Declaration of generative AI and AI-assisted technologies

During the preparation of this work, the author(s) used ChatGPT to refine the text and code. After using this tool or service, the author(s) reviewed and edited the content as needed and take(s) full responsibility for the content of the publication.

