## Supplemental Figures for "Multi-omic landscape of human gliomas from diagnosis to treatment and recurrence"

A

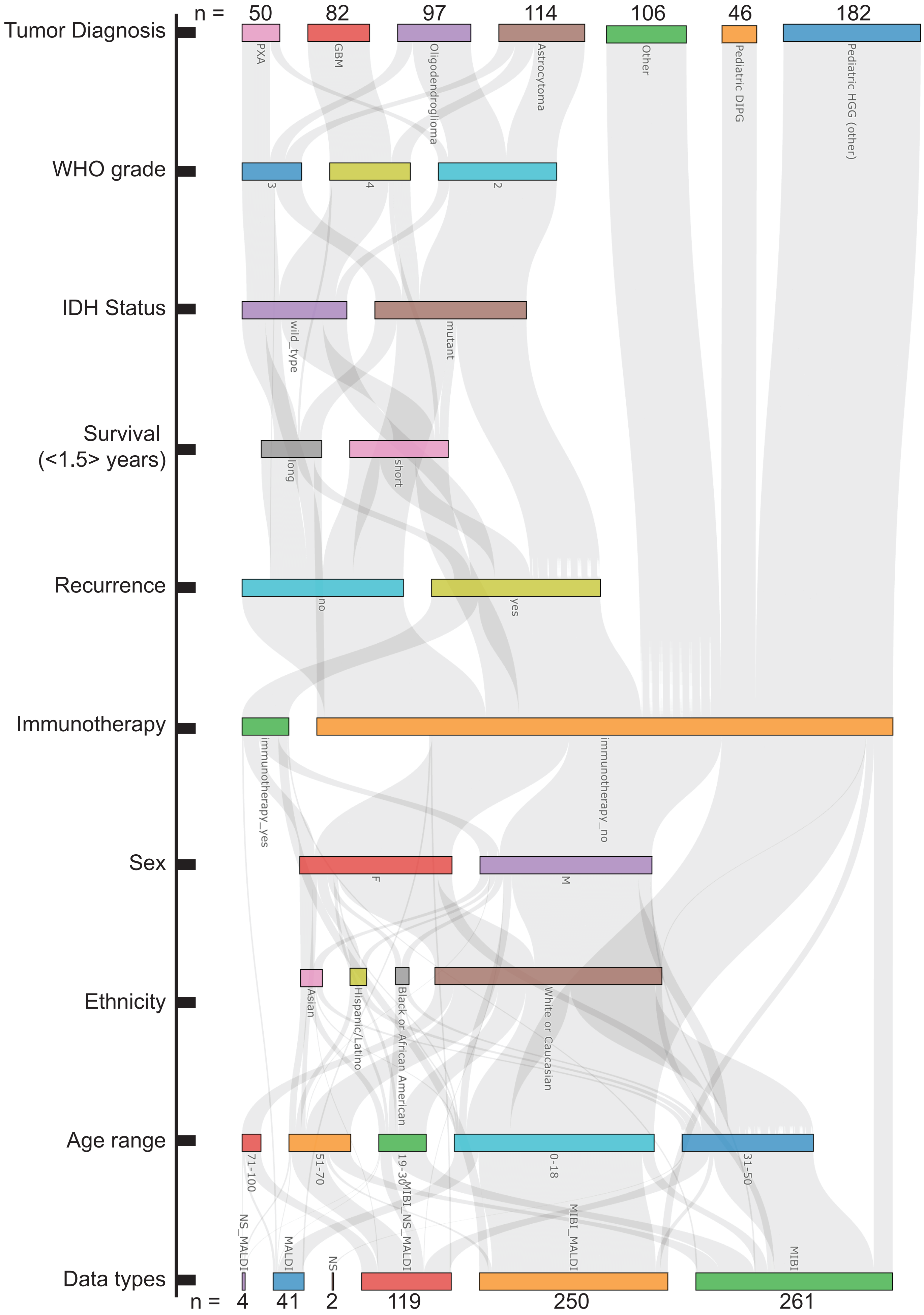
n =
50
82
97
114
106
46
182
Tumor Diagnosis
PXA
GBM
Oligodendroglioma
Astrocytoma
Other
Pediatric DIPG
Pediatric HGG (other)
WHO grade
3
4
2
IDH Status
wild_type
mutant
Survival
(<1.5> years)
long
short
Recurrence
no
yes
Immunotherapy
immunotherapy_yes
immunotherapy_no
Sex
F
M
Ethnicity
Asian
Hispanic/Latino
Black or African American
White or Caucasian
Age range
71-100
51-70
19-30
0-18
31-50
Data types
NS_MALDI
MALDI
NS
MIBI_NS_MALDI
MIBI_MALDI
MIBI
n =
4
41
2
119
250
261

**Figure S1. Metadata and data type of BRUCE cohort, related to Figure 1.**
**(A)** Bruce cohort samples identified by metadata and data type (Tumor Diagnosis, WHO Grade, IDH Status, Survival (<1.5> years), Recurrence, Immunotherapy, Sex, Ethnicity, Age Range, and Data Type).

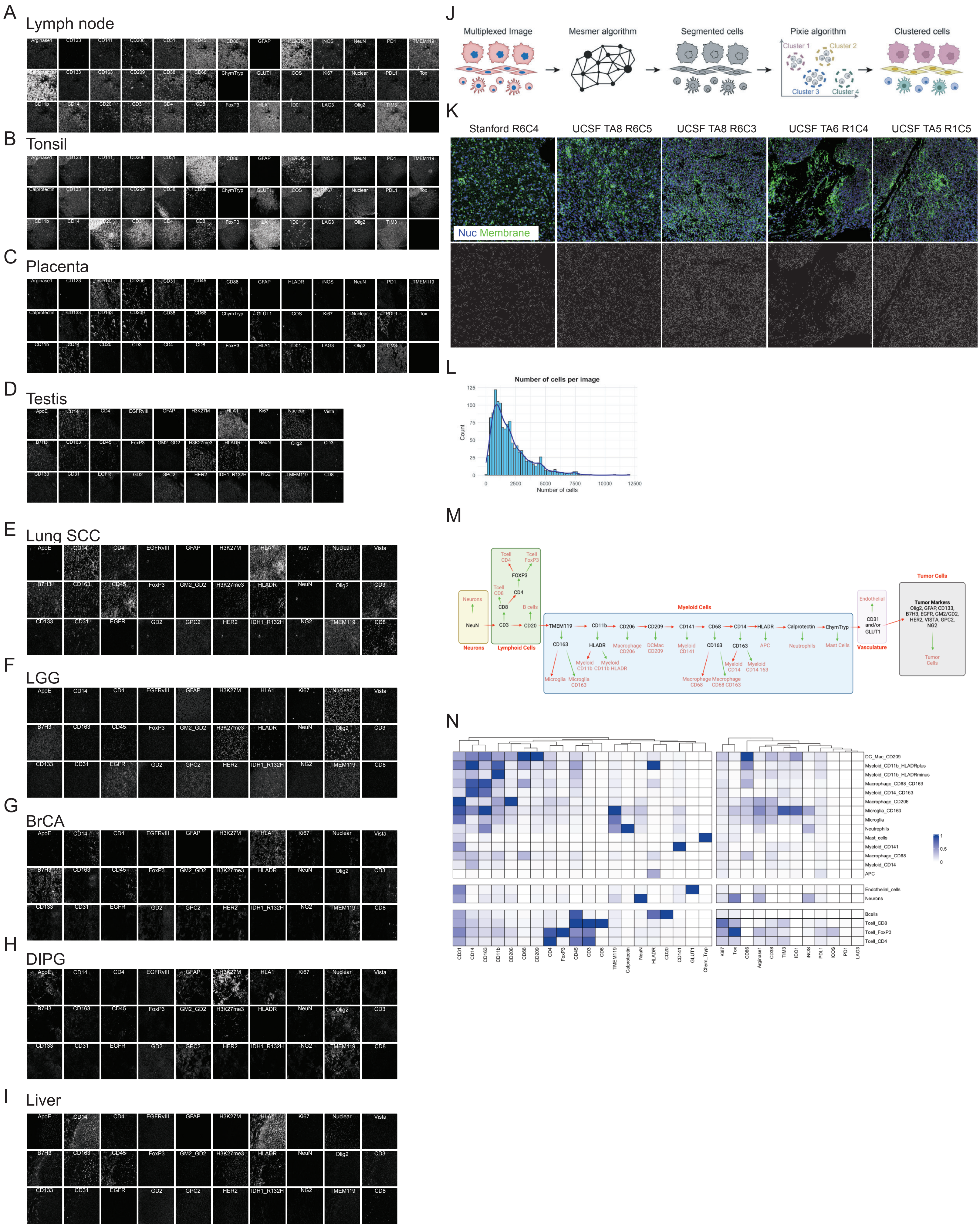

A
Lymph node
B
Tonsil
C
Placenta
D
Testis
E
Lung SCC
F
LGG
G
BrCA
H
DIPG
I
Liver
J
Multiplexed Image
Mesmer algorithm
Segmented cells
Pixie algorithm
Cluster 1
Cluster 2
Cluster 3
Cluster 4
Clustered cells
K
Stanford R6C4
UCSF TA8 R6C5
UCSF TA8 R6C3
UCSF TA6 R1C4
UCSF TA5 R1C5
Nuc Membrane
L
Number of cells per image
Count
Number of cells
M
Neurons
NeuN
Lymphoid Cells
Myeloid Cells
Vasculature
Tumor Cells
Tumor Markers
Olig2, GFAP, CD133, B7H3, EGFR, GM2/GD2, HER2, VISTA, GPC2, NG2
N
DC_Mac_CD209
Myeloid_CD11b_HLADRplus
Myeloid_CD11b_HLADRminus
Macrophage_CD68_CD163
Myeloid_CD14_CD163
Macrophage_CD206
Microglia_CD163
Microglia
Neutrophils
Mast_cells
Myeloid_CD141
Macrophage_CD68
Myeloid_CD14
APC
Endothelial_cells
Neurons
Bcells
Tcell_CD8
Tcell_FoxP3
Tcell_CD4

**Figure S2. Tissue QC and preprocessing and validation, related to Figures 1 and 2.**
**(A-I)** Grayscale images of marker signal on control tissues (lymph node, tonsil, placenta, testis, lung squamous carcinoma, low grade glioma, breast cancer, diffuse intrinsic pontine glioma, liver) **(J)** Schematic showing the steps of the Mesmer pipeline. **(K)** MIBI images show nuclear (blue) and membrane (green) proteins with associated whole cell segmentation (top) and individual identified and segmented cells (bottom). **(L)** Histogram showing the frequency of images with varying cell counts, while the overlaid density curve represents the smoothed probability distribution. **(M)** Representative schematic of the thresholding strategy used to phenotype cells. **(N)** Immune cell lineage assignments based on normalized expression of lineage markers. Rows have been grouped by cell subtype (myeloid, endothelial/neuron, and lymphoid) columns are hierarchically clustered (Euclidean distance, average linkage). Z scored by row.

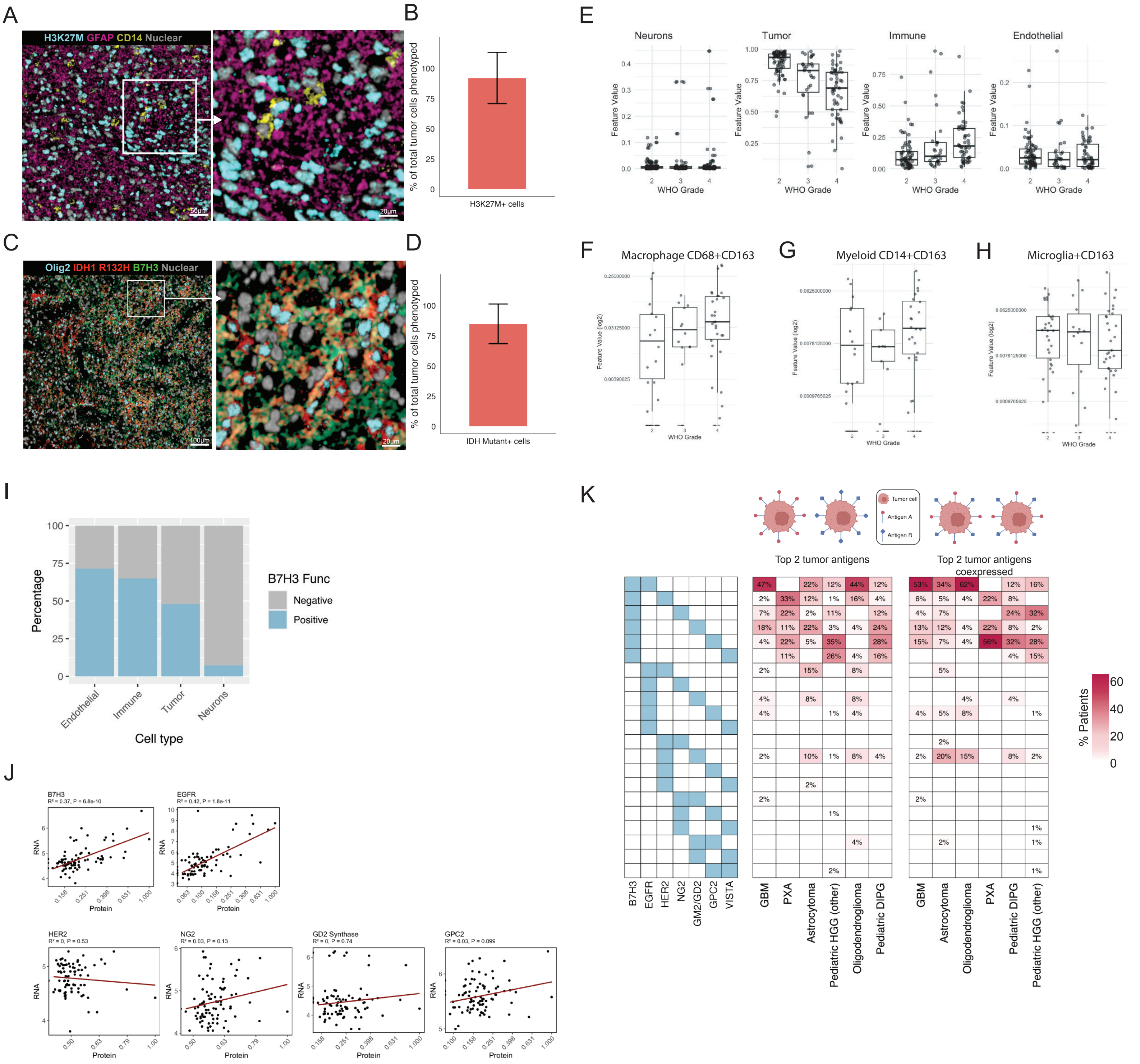
A
H3K27M GFAP CD14 Nuclear
50μm
20μm
B
% of total tumor cells phenotyped
H3K27M+ cells
C
Olig2 IDH1 R132H B7H3 Nuclear
100μm
20μm
D
% of total tumor cells phenotyped
IDH Mutant+ cells
E
Neurons
Tumor
Immune
Endothelial
Feature Value
WHO Grade
F
Macrophage CD68+CD163
G
Myeloid CD14+CD163
H
Microglia+CD163
Feature Value (log2)
WHO Grade
I
Percentage
B7H3 Func
Negative
Positive
Endothelial
Immune
Tumor
Neurons
Cell type
J
B7H3
R² = 0.37, P = 6.8e-10
EGFR
R² = 0.42, P = 1.8e-11
HER2
R² = 0, P = 0.53
NG2
R² = 0.03, P = 0.13
GD2 Synthase
R² = 0, P = 0.74
GPC2
R² = 0.03, P = 0.099
RNA
Protein
K
Tumor cell
Antigen A
Antigen B
Top 2 tumor antigens
Top 2 tumor antigens coexpressed
B7H3
EGFR
HER2
NG2
GM2/GD2
GPC2
VISTA
GBM
PXA
Astrocytoma
Pediatric HGG (other)
Oligodendroglioma
Pediatric DIPG
% Patients

**Figure S3. Immune and tumor antigen metrics, related to Figures 2 and 3. (A, C)** Representative multiplexed image. **(B)** Bar graph represents the percentage of total tumor cells that are H3K27M+. **(D)** Bar graph represents the percentage of total tumor cells that are IDH1 R132H+. **(E)** Boxplot showing the per patient percentage of total cells that have been identified as Neuron, Tumor, Immune, or Endothelial stratified by WHO grade. **(F-H)** % of different CD163+ immune cells relative to all immune cells by WHO grade. **()** % of different cell types colored by B7H3 positivity **(J)** Top two tumor antigen expression pattern heatmap shows % patients for combinations of top 2 TA yielding maximal coverage of tumor cells for each glioma subtype (left). Heatmap shows % patients for combinations of top 2 TA that are co-expressed on the same tumor cell for each glioma subtype (right). **(K)** RNA and protein correlation plot for TA.

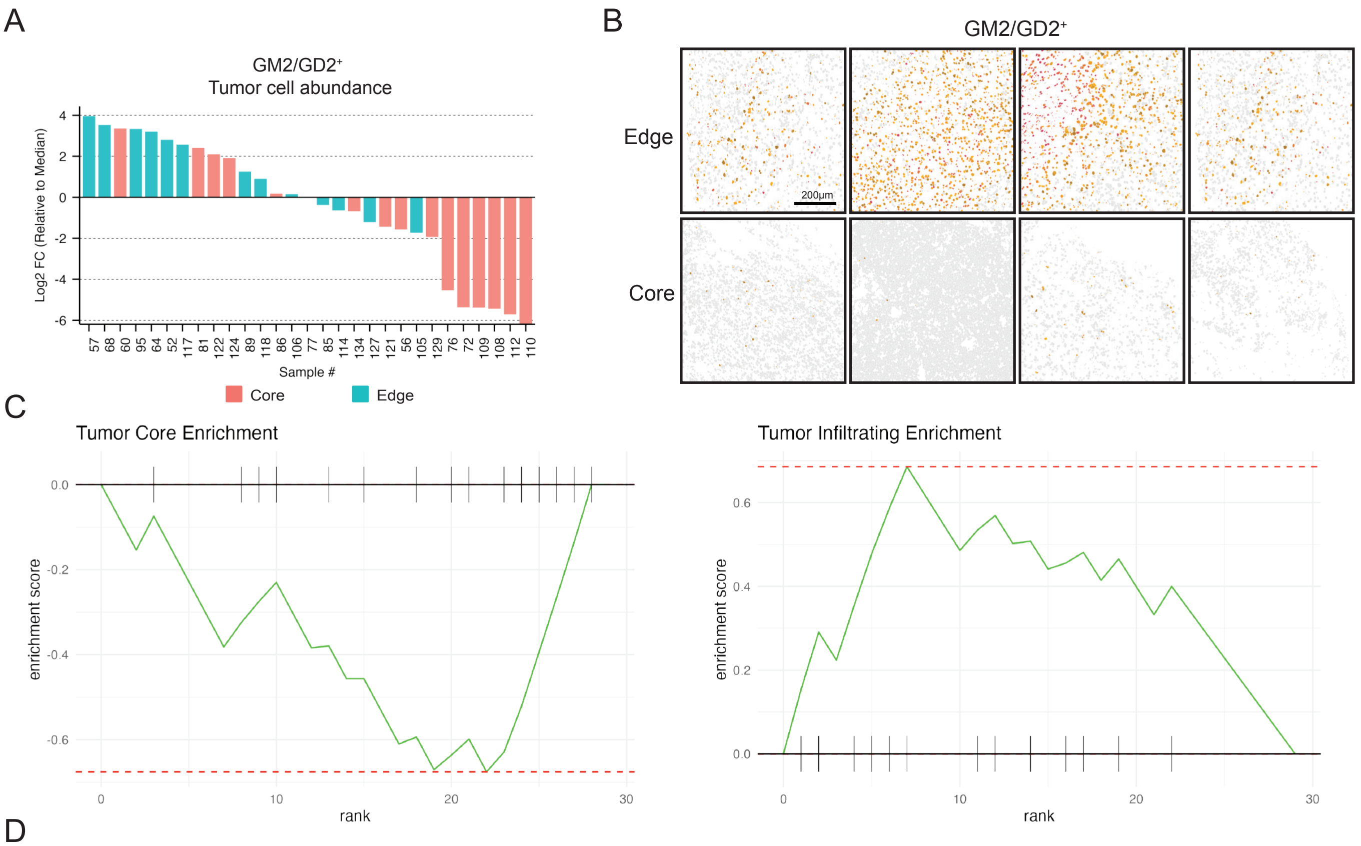


D

| pathway | pval | padj | log2err | ES | NES | size | leadingEdge |
|---|---|---|---|---|---|---|---|
| Tumor_Core | 0.014048289 | 0.014048289 | 0.380730401 | -0.675802071 | -1.686951259 | 15 | 112, 108, 109, 72, 76, 129 |
| Tumor_Infiltrating | 0.009601796 | 0.014048289 | 0.380730401 | 0.685807296 | 1.861938813 | 13 | 57, 68, 95, 64, 52, 117 |

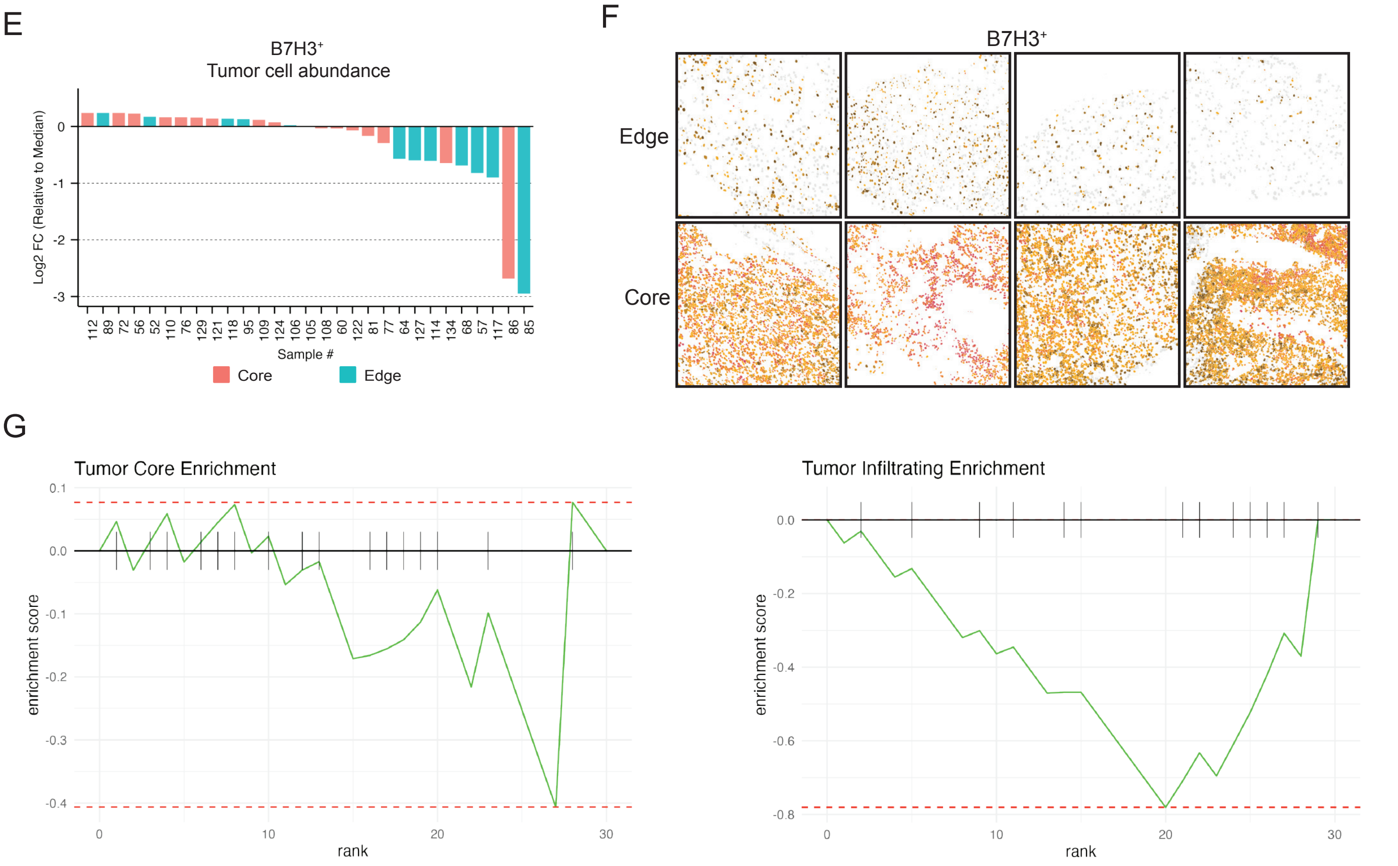


H

| pathway | pval | padj | log2err | ES | NES | size | leadingEdge |
|---|---|---|---|---|---|---|---|
| Tumor_Core | 0.893305439 | 0.893305439 | 0.018928789 | -0.406263953 | -0.702075147 | 16 | 86 |
| Tumor_Infiltrating | 0.065887354 | 0.131774708 | 0.178219875 | -0.780499008 | -1.351094002 | 13 | 85, 117, 57, 127, 68, 114, 64 |

**Figure S4. B7H3 and GM2/GD2 abundance between the core and infiltrating edge, related to Figure 4. (A)** Bar plot (each bar shows unique sample) showing the log2 fold change of GM2/GD2+ tumor cell abundance relative to the median across all treatment naive GBM samples, colored by tumor core (blue, filled) and edge (blue, unfilled) regions. **(B)** Spatial distribution of GM2/GD2+ tumor cell abundance in representative tumor sections, showing the tumor edge and core regions. Color represents antigen intensity (High - red, Medium - orange, Low - black, Negative - grey) Scale bar = 200 μm. **(C)** Gene set enrichment analysis (GSEA) on GM2/GD2+ bar plots in which each "gene" is a patient sample and each "gene set" is a tumor region. Dotted line represents either the minimum or maximum enrichment score. **(D)** Resulting gene set analysis statistics. **(E)** Bar plot (each bar shows unique sample) showing the log2 fold change of B7H3+ tumor cell abundance relative to the median across all treatment naive GBM samples, colored by tumor core (blue, filled) and edge (blue, unfilled) regions. **(F)** Spatial distribution of B7H3+ tumor cell abundance in representative tumor sections, showing the tumor edge and core regions. Color represents antigen intensity (High - red, Medium - orange, Low - black, Negative - grey) **(G)** Gene set enrichment analysis on B7H3+ bar plots in which each "gene" is a patient sample and each "gene set" is a tumor region. Dotted line represents either the minimum or maximum enrichment score. **(H)** Resulting gene set analysis statistics.

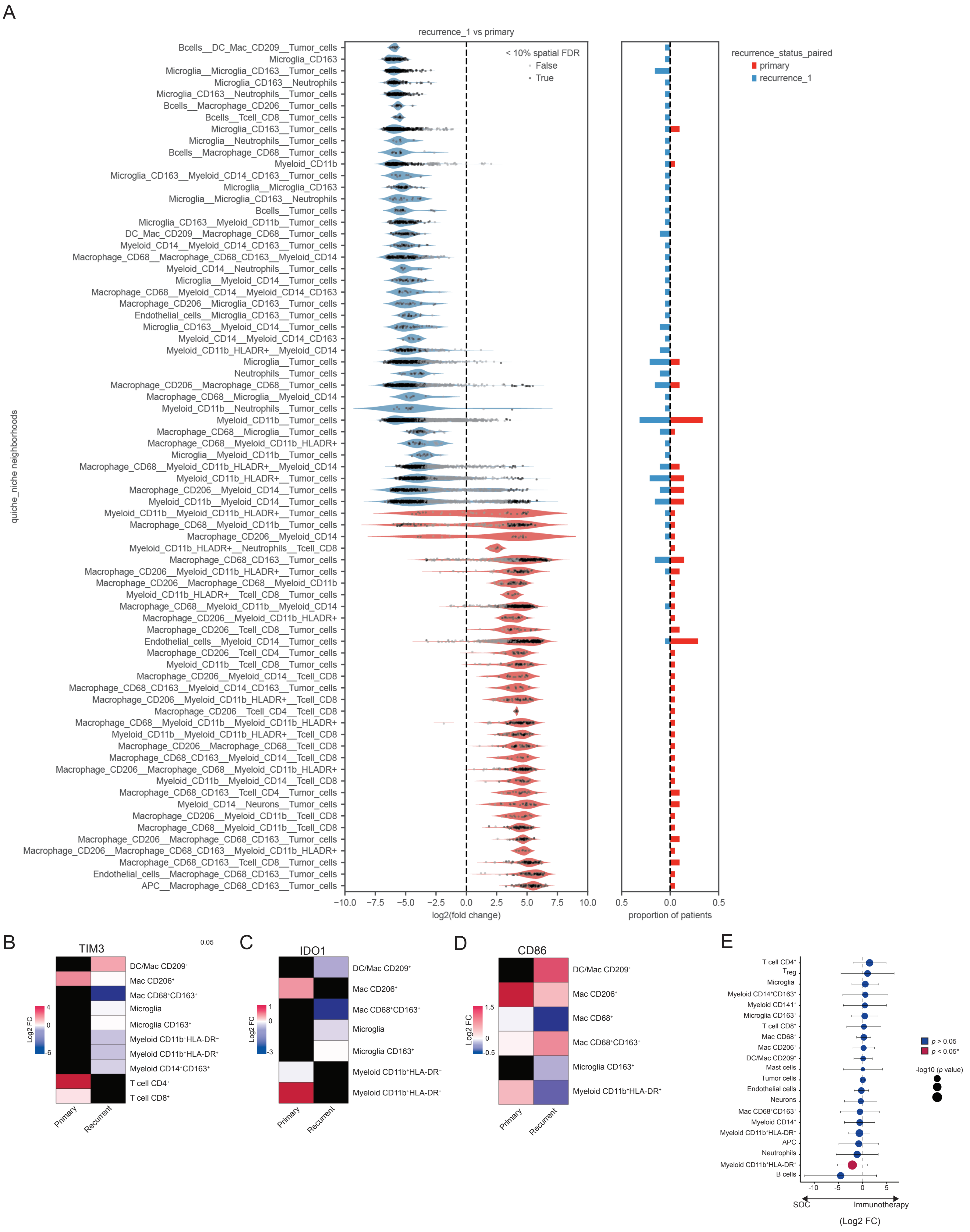
A
recurrence_1 vs primary
< 10% spatial FDR
False
True
recurrence_status_paired
primary
recurrence_1
quiche_niche neighborhoods
Bcells__DC_Mac_CD209__Tumor_cells
Microglia_CD163
Microglia__Microglia_CD163__Tumor_cells
Microglia_CD163__Neutrophils
Microglia_CD163__Neutrophils__Tumor_cells
Bcells__Macrophage_CD206__Tumor_cells
Bcells__Tcell_CD8__Tumor_cells
Microglia_CD163__Tumor_cells
Microglia__Neutrophils__Tumor_cells
Bcells__Macrophage_CD68__Tumor_cells
Myeloid_CD11b
Microglia_CD163__Myeloid_CD14_CD163__Tumor_cells
Microglia__Microglia_CD163
Microglia__Microglia_CD163__Neutrophils
Bcells__Tumor_cells
Microglia_CD163__Myeloid_CD11b__Tumor_cells
DC_Mac_CD209__Macrophage_CD68__Tumor_cells
Myeloid_CD14__Myeloid_CD14_CD163__Tumor_cells
Macrophage_CD68__Macrophage_CD68_CD163__Myeloid_CD14
Myeloid_CD14__Neutrophils__Tumor_cells
Microglia__Myeloid_CD14__Tumor_cells
Macrophage_CD68__Myeloid_CD14__Myeloid_CD14_CD163
Macrophage_CD206__Microglia_CD163__Tumor_cells
Endothelial_cells__Microglia_CD163__Tumor_cells
Microglia_CD163__Myeloid_CD14__Tumor_cells
Myeloid_CD14__Myeloid_CD14_CD163
Myeloid_CD11b_HLADR+__Myeloid_CD14
Microglia__Tumor_cells
Neutrophils__Tumor_cells
Macrophage_CD206__Macrophage_CD68__Tumor_cells
Macrophage_CD68__Microglia__Myeloid_CD14
Myeloid_CD11b__Neutrophils__Tumor_cells
Myeloid_CD11b__Tumor_cells
Macrophage_CD68__Microglia__Tumor_cells
Macrophage_CD68__Myeloid_CD11b_HLADR+
Microglia__Myeloid_CD11b__Tumor_cells
Macrophage_CD68__Myeloid_CD11b_HLADR+__Myeloid_CD14
Myeloid_CD11b_HLADR+__Tumor_cells
Macrophage_CD206__Myeloid_CD14__Tumor_cells
Myeloid_CD11b__Myeloid_CD14__Tumor_cells
Myeloid_CD11b__Myeloid_CD11b_HLADR+__Tumor_cells
Macrophage_CD68__Myeloid_CD11b__Tumor_cells
Macrophage_CD206__Myeloid_CD14
Myeloid_CD11b_HLADR+__Neutrophils__Tcell_CD8
Macrophage_CD68_CD163__Tumor_cells
Macrophage_CD206__Myeloid_CD11b_HLADR+__Tumor_cells
Macrophage_CD206__Macrophage_CD68__Myeloid_CD11b
Myeloid_CD11b_HLADR+__Tcell_CD8__Tumor_cells
Macrophage_CD68__Myeloid_CD11b__Myeloid_CD14
Macrophage_CD206__Myeloid_CD11b_HLADR+
Macrophage_CD206__Tcell_CD8__Tumor_cells
Endothelial_cells__Myeloid_CD14__Tumor_cells
Macrophage_CD206__Tcell_CD4__Tumor_cells
Myeloid_CD11b__Tcell_CD8__Tumor_cells
Macrophage_CD206__Myeloid_CD14__Tcell_CD8
Macrophage_CD68_CD163__Myeloid_CD14_CD163__Tumor_cells
Macrophage_CD206__Myeloid_CD11b_HLADR+__Tcell_CD8
Macrophage_CD206__Tcell_CD4__Tcell_CD8
Macrophage_CD68__Myeloid_CD11b__Myeloid_CD11b_HLADR+
Myeloid_CD11b__Myeloid_CD11b_HLADR+__Tcell_CD8
Macrophage_CD206__Macrophage_CD68__Tcell_CD8
Macrophage_CD68_CD163__Myeloid_CD14__Tcell_CD8
Macrophage_CD206__Macrophage_CD68__Myeloid_CD11b_HLADR+
Myeloid_CD11b__Myeloid_CD14__Tcell_CD8
Macrophage_CD68_CD163__Tcell_CD4__Tumor_cells
Myeloid_CD14__Neurons__Tumor_cells
Macrophage_CD206__Myeloid_CD11b__Tcell_CD8
Macrophage_CD68__Myeloid_CD11b__Tcell_CD8
Macrophage_CD206__Macrophage_CD68_CD163__Tumor_cells
Macrophage_CD206__Macrophage_CD68_CD163__Myeloid_CD11b_HLADR+
Macrophage_CD68_CD163__Tcell_CD8__Tumor_cells
Endothelial_cells__Macrophage_CD68_CD163__Tumor_cells
APC__Macrophage_CD68_CD163__Tumor_cells
−10.0 −7.5 −5.0 −2.5 0.0 2.5 5.0 7.5 10.0
log2(fold change)
0.5 0.0 0.5
proportion of patients
B
TIM3
0.05
DC/Mac CD209+
Mac CD206+
Mac CD68+CD163+
Microglia
Microglia CD163+
Myeloid CD11b+HLA-DR−
Myeloid CD11b+HLA-DR+
Myeloid CD14+CD163+
T cell CD4+
T cell CD8+
Log2 FC
4
0
-6
Primary
Recurrent
C
IDO1
DC/Mac CD209+
Mac CD206+
Mac CD68+CD163+
Microglia
Microglia CD163+
Myeloid CD11b+HLA-DR−
Myeloid CD11b+HLA-DR+
Log2 FC
1
0
-3
Primary
Recurrent
D
CD86
DC/Mac CD209+
Mac CD206+
Mac CD68+
Mac CD68+CD163+
Microglia CD163+
Myeloid CD11b+HLA-DR+
Log2 FC
1.5
0
-0.5
Primary
Recurrent
E
T cell CD4+
Treg
Microglia
Myeloid CD14+CD163+
Myeloid CD141+
Microglia CD163+
T cell CD8+
Mac CD68+
Mac CD206+
DC/Mac CD209+
Mast cells
Tumor cells
Endothelial cells
Neurons
Mac CD68+CD163+
Myeloid CD14+
Myeloid CD11b+HLA-DR−
APC
Neutrophils
Myeloid CD11b+HLA-DR+
B cells
-10 -5 0 5
SOC
Immunotherapy
(Log2 FC)
p > 0.05
p < 0.05*
-log10 (p value)

**Figure S5. QUICHE analysis details, related to Figure 5. (A)** Violin plots show the top differentially abundant niche neighborhoods in patients who are recurrent (blue) or primary (red). Bar plots to the right show the proportion of patients with a niche neighborhood in the respective patient groups. **(B-D)** Functional marker analysis shows differential enrichment of TIM3, IDO1, and CD86 (Log2 fold change between differentially abundant niche neighborhoods to broader TME) in primary and recurrent tumors. **(E)** Differential abundance analysis of immune cell types between SOC and immunotherapy groups, showing log2 fold changes and statistical significance (-log10 p-value).

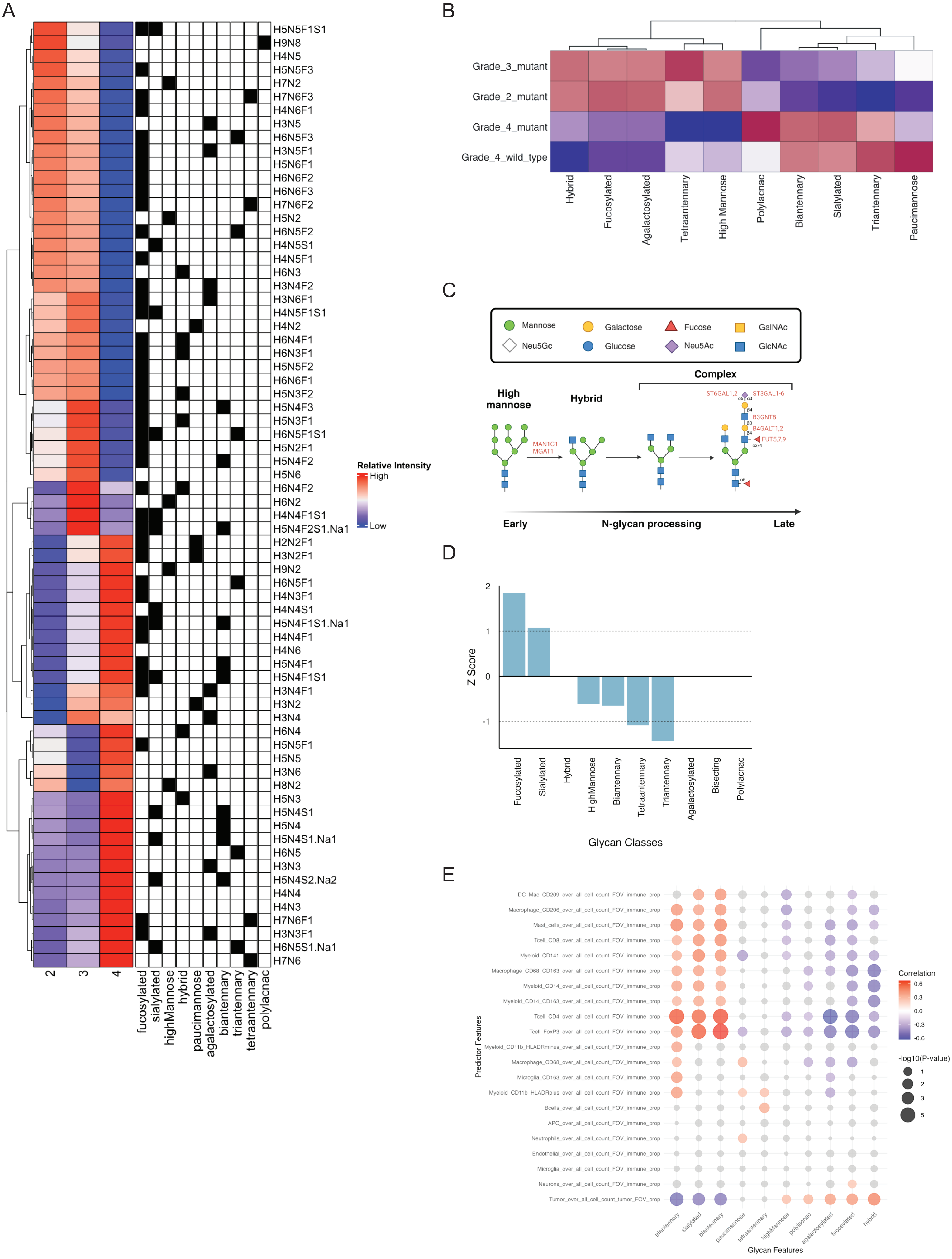
A
H5N5F1S1
H9N8
H4N5
H5N5F3
H7N2
H7N6F3
H4N6F1
H3N5
H6N5F3
H3N5F1
H5N6F1
H6N6F2
H6N6F3
H7N6F2
H5N2
H6N5F2
H4N5S1
H4N5F1
H6N3
H3N4F2
H3N6F1
H4N5F1S1
H4N2
H6N4F1
H6N3F1
H5N5F2
H6N6F1
H5N3F2
H5N4F3
H5N3F1
H6N5F1S1
H5N2F1
H5N4F2
H5N6
H6N4F2
H6N2
H4N4F1S1
H5N4F2S1.Na1
H2N2F1
H3N2F1
H9N2
H6N5F1
H4N3F1
H4N4S1
H5N4F1S1.Na1
H4N4F1
H4N6
H5N4F1
H5N4F1S1
H3N4F1
H3N2
H3N4
H6N4
H5N5F1
H5N5
H3N6
H8N2
H5N3
H5N4S1
H5N4
H5N4S1.Na1
H6N5
H3N3
H5N4S2.Na2
H4N4
H4N3
H7N6F1
H3N3F1
H6N5S1.Na1
H7N6
2
3
4
fucosylated
sialylated
highMannose
hybrid
paucimannose
agalactosylated
biantennary
triantennary
tetraantennary
polylacnac
Relative Intensity
High
Low
B
Grade_3_mutant
Grade_2_mutant
Grade_4_mutant
Grade_4_wild_type
Hybrid
Fucosylated
Agalactosylated
Tetraantennary
High Mannose
Polylacnac
Biantennary
Sialylated
Triantennary
Paucimannose
C
Mannose
Galactose
Fucose
GalNAc
Neu5Gc
Glucose
Neu5Ac
GlcNAc
Complex
High mannose
Hybrid
MAN1C1
MGAT1
ST6GAL1,2
ST3GAL1-6
B3GNT8
B4GALT1,2
FUT5,7,9
Early
N-glycan processing
Late
D
2
1
0
-1
Z Score
Fucosylated
Sialylated
Hybrid
HighMannose
Biantennary
Tetraantennary
Triantennary
Agalactosylated
Bisecting
Polylacnac
Glycan Classes
E
DC_Mac_CD209_over_all_cell_count_FOV_immune_prop
Macrophage_CD206_over_all_cell_count_FOV_immune_prop
Mast_cells_over_all_cell_count_FOV_immune_prop
Tcell_CD8_over_all_cell_count_FOV_immune_prop
Myeloid_CD141_over_all_cell_count_FOV_immune_prop
Macrophage_CD68_CD163_over_all_cell_count_FOV_immune_prop
Myeloid_CD14_over_all_cell_count_FOV_immune_prop
Myeloid_CD14_CD163_over_all_cell_count_FOV_immune_prop
Tcell_CD4_over_all_cell_count_FOV_immune_prop
Tcell_FoxP3_over_all_cell_count_FOV_immune_prop
Myeloid_CD11b_HLADRminus_over_all_cell_count_FOV_immune_prop
Macrophage_CD68_over_all_cell_count_FOV_immune_prop
Microglia_CD163_over_all_cell_count_FOV_immune_prop
Myeloid_CD11b_HLADRplus_over_all_cell_count_FOV_immune_prop
Bcells_over_all_cell_count_FOV_immune_prop
APC_over_all_cell_count_FOV_immune_prop
Neutrophils_over_all_cell_count_FOV_immune_prop
Endothelial_over_all_cell_count_FOV_immune_prop
Microglia_over_all_cell_count_FOV_immune_prop
Neurons_over_all_cell_count_FOV_immune_prop
Tumor_over_all_cell_count_tumor_FOV_immune_prop
Predictor Features
triantennary
sialylated
biantennary
paucimannose
tetraantennary
highMannose
polylacnac
agalactosylated
fucosylated
hybrid
Glycan Features
Correlation
0.6
0.3
0.0
-0.3
-0.6
-log10(P-value)
1
2
3
5

**Figure S6. Glycan expression, related to Figure 6. (A)** Heatmap showing differences in relative intensity of all 70 glycans identified by WHO grade. **(B)** Heatmap showing differences in relative intensity of all 70 glycans identified by WHO grade and IDH status. **(C)** Schematic representation of N-glycan processing, from early-stage high mannose glycans to more complex structures, including hybrid and complex glycans. The diagram highlights key enzymes involved. **(D)** Z-score analysis of the co-occurrence of glycan classes and the enrichment of enzymes related to the processing of the glycan class. Positive Z-score > 1 identifies the coordination is more than random and Z-score < -1 shows coordination less than random. **(E)** Correlation and p value of cell types to glycan classes.

A

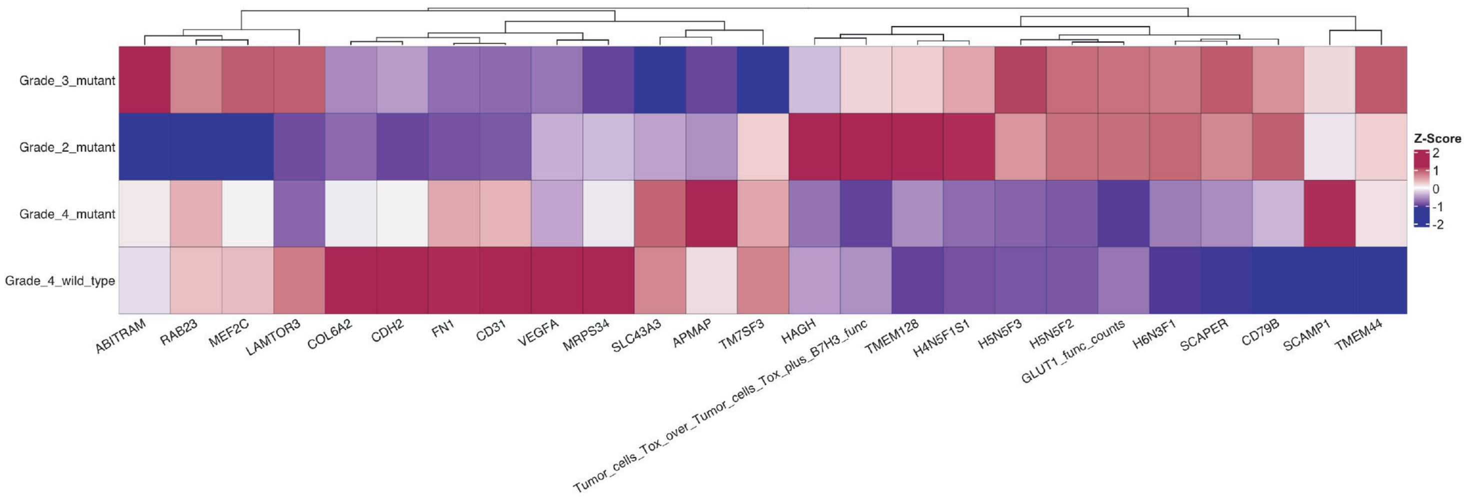
Grade_3_mutant
Grade_2_mutant
Grade_4_mutant
Grade_4_wild_type
Z-Score
2
1
0
-1
-2
ABITRAM
RAB23
MEF2C
LAMTOR3
COL6A2
CDH2
FN1
CD31
VEGFA
MRPS34
SLC43A3
APMAP
TM7SF3
HAGH
Tumor_cells_Tox_over_Tumor_cells_Tox_plus_B7H3_func
TMEM128
H4N5F1S1
H5N5F3
H5N5F2
GLUT1_func_counts
H6N3F1
SCAPER
CD79B
SCAMP1
TMEM44

B

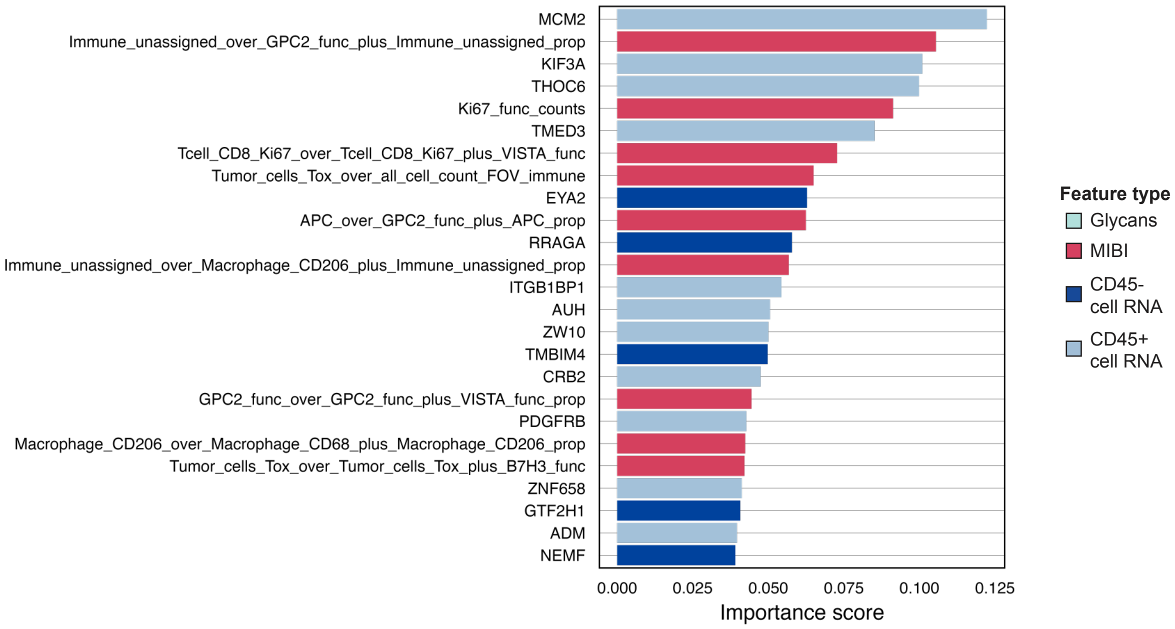
MCM2
Immune_unassigned_over_GPC2_func_plus_Immune_unassigned_prop
KIF3A
THOC6
Ki67_func_counts
TMED3
Tcell_CD8_Ki67_over_Tcell_CD8_Ki67_plus_VISTA_func
Tumor_cells_Tox_over_all_cell_count_FOV_immune
EYA2
APC_over_GPC2_func_plus_APC_prop
RRAGA
Immune_unassigned_over_Macrophage_CD206_plus_Immune_unassigned_prop
ITGB1BP1
AUH
ZW10
TMBIM4
CRB2
GPC2_func_over_GPC2_func_plus_VISTA_func_prop
PDGFRB
Macrophage_CD206_over_Macrophage_CD68_plus_Macrophage_CD206_prop
Tumor_cells_Tox_over_Tumor_cells_Tox_plus_B7H3_func
ZNF658
GTF2H1
ADM
NEMF
0.000
0.025
0.050
0.075
0.100
0.125
Importance score
Feature type
Glycans
MIBI
CD45-
cell RNA
CD45+
cell RNA

**Figure S7. Classifier details, related to Figure 7. (A)** Heatmap showing median importance of top 25 features for WHO grade classifier in IDH status **(B)** Top 25 features for survival classifier.

### Table S1. Patient data

| Features | Values | CHOP[a] | CoH[b] | Stanford | UCLA[c] | UCSF[d] |
|---|---|---|---|---|---|---|
| Disease | Breast Cancer | 4 (2.56%) | 0 (0%) | 0 (0%) | 0 (0%) | 0 (0%) |
| | Colon Cancer | 4 (2.56%) | 0 (0%) | 0 (0%) | 0 (0%) | 0 (0%) |
| | Placenta | 24 (15.38%) | 0 (0%) | 1 (1.16%) | 0 (0%) | 0 (0%) |
| | Tonsil | 6 (3.85%) | 0 (0%) | 0 (0%) | 0 (0%) | 0 (0%) |
| | Pediatric Astrocytoma | 24 (15.38%) | 0 (0%) | 0 (0%) | 0 (0%) | 0 (0%) |
| | Pediatric Diffuse Midline Glioma | 25 (16.03%) | 0 (0%) | 0 (0%) | 0 (0%) | 0 (0%) |
| | Pediatric GBM[e] | 33 (21.15%) | 0 (0%) | 0 (0%) | 0 (0%) | 0 (0%) |
| | Pediatric Ganglioglioma | 2 (1.28%) | 0 (0%) | 0 (0%) | 0 (0%) | 0 (0%) |
| | Pediatric Glioma | 16 (10.26%) | 0 (0%) | 0 (0%) | 0 (0%) | 0 (0%) |
| | Pediatric HGG[f] | 8 (5.13%) | 0 (0%) | 0 (0%) | 0 (0%) | 0 (0%) |
| | Pediatric PXA[g] | 8 (5.13%) | 0 (0%) | 0 (0%) | 0 (0%) | 0 (0%) |
| | Pediatric Thalmic Glioma | 2 (1.28%) | 0 (0%) | 0 (0%) | 0 (0%) | 0 (0%) |
| | GBM | 0 (0%) | 6 (100%) | 32 (37.21%) | 12 (100%) | 0 (0%) |
| | Astrocytoma | 0 (0%) | 0 (0%) | 22 (25.58%) | 0 (0%) | 20 (37.74%) |
| | FCD[h] | 0 (0%) | 0 (0%) | 1 (1.16%) | 0 (0%) | 0 (0%) |
| | FTC[i] | 0 (0%) | 0 (0%) | 8 (9.3%) | 0 (0%) | 0 (0%) |
| | GBM Other | 0 (0%) | 0 (0%) | 1 (1.16%) | 0 (0%) | 0 (0%) |
| | Normal Brain | 0 (0%) | 0 (0%) | 1 (1.16%) | 0 (0%) | 4 (7.55%) |
| | Oligodendroglioma | 0 (0%) | 0 (0%) | 8 (9.3%) | 0 (0%) | 19 (35.85%) |
| | VM[j] | 0 (0%) | 0 (0%) | 3 (3.49%) | 0 (0%) | 0 (0%) |
| | gliosis | 0 (0%) | 0 (0%) | 7 (8.14%) | 0 (0%) | 0 (0%) |
| | mFTC[k] brain | 0 (0%) | 0 (0%) | 2 (2.33%) | 0 (0%) | 0 (0%) |
| | Meningioma | 0 (0%) | 0 (0%) | 0 (0%) | 0 (0%) | 1 (1.89%) |
| | PXA | 0 (0%) | 0 (0%) | 0 (0%) | 0 (0%) | 9 (16.98%) |
| Sex | Female | 19 (12.18%) | 3 (50%) | 37 (43.02%) | 0 (0%) | 19 (35.85%) |
| | Male | 15 (9.62%) | 3 (50%) | 45 (52.33%) | 0 (0%) | 29 (54.72%) |
| | Unknown | 122 (78.21%) | 0 (0%) | 4 (4.65%) | 12 (100%) | 5 (9.43%) |
| Race | Asian | 1 (0.64%) | 0 (0%) | 5 (5.81%) | 0 (0%) | 3 (5.66%) |
| | Black or African American | 1 (0.64%) | 0 (0%) | 0 (0%) | 0 (0%) | 3 (5.66%) |
| | White or Caucasian | 27 (17.31%) | 0 (0%) | 56 (65.12%) | 0 (0%) | 35 (66.04%) |
| | unknown | 127 (81.41%) | 6 (100%) | 7 (8.14%) | 12 (100%) | 12 (22.64%) |
| | Hispanic/Latino | 0 (0%) | 0 (0%) | 15 (17.44%) | 0 (0%) | 0 (0%) |
| | NA | 0 (0%) | 0 (0%) | 3 (3.49%) | 0 (0%) | 0 (0%) |
| Immunotherapy | no | 156 (100%) | 3 (50%) | 86 (100%) | 4 (33.33%) | 35 (66.04%) |
| | yes | 0 (0%) | 3 (50%) | 0 (0%) | 8 (66.67%) | 18 (33.96%) |
| WHO Grade | NA | 156 (100%) | 0 (0%) | 24 (27.91%) | 0 (0%) | 5 (9.43%) |
| | 4 | 0 (0%) | 6 (100%) | 40 (46.51%) | 12 (100%) | 2 (3.77%) |
| | 2 | 0 (0%) | 0 (0%) | 10 (11.63%) | 0 (0%) | 38 (71.7%) |
| | 3 | 0 (0%) | 0 (0%) | 12 (13.95%) | 0 (0%) | 8 (15.09%) |
| IDH Status | unknown | 156 (100%) | 0 (0%) | 27 (31.4%) | 0 (0%) | 5 (9.43%) |
| | wild type | 0 (0%) | 6 (100%) | 36 (41.86%) | 12 (100%) | 9 (16.98%) |
| | mutant | 0 (0%) | 0 (0%) | 23 (26.74%) | 0 (0%) | 39 (73.58%) |

| | | | | | | |
|---|---|---|---|---|---|---|
| MGMT Status | NA | 156 (100%) | 6 (100%) | 41 (47.67%) | 10 (83.33%) | 53 (100%) |
| | mutant | 0 (0%) | 0 (0%) | 24 (27.91%) | 2 (16.67%) | 0 (0%) |
| | wild type | 0 (0%) | 0 (0%) | 21 (24.42%) | 0 (0%) | 0 (0%) |
| P53 Status | NA | 156 (100%) | 6 (100%) | 47 (54.65%) | 12 (100%) | 21 (39.62%) |
| | Stain retained | 0 (0%) | 0 (0%) | 2 (2.33%) | 0 (0%) | 0 (0%) |
| | mutant | 0 (0%) | 0 (0%) | 16 (18.6%) | 0 (0%) | 15 (28.3%) |
| | wild type | 0 (0%) | 0 (0%) | 21 (24.42%) | 0 (0%) | 17 (32.08%) |
| Tertpro Status | NA | 156 (100%) | 6 (100%) | 86 (100%) | 12 (100%) | 51 (96.23%) |
| | 1 | 0 (0%) | 0 (0%) | 0 (0%) | 0 (0%) | 2 (3.77%) |
| ATRX Status | NA | 156 (100%) | 6 (100%) | 52 (60.47%) | 12 (100%) | 23 (43.4%) |
| | Stain retained | 0 (0%) | 0 (0%) | 1 (1.16%) | 0 (0%) | 0 (0%) |
| | conserved | 0 (0%) | 0 (0%) | 1 (1.16%) | 0 (0%) | 0 (0%) |
| | mutant | 0 (0%) | 0 (0%) | 3 (3.49%) | 0 (0%) | 17 (32.08%) |
| | mutant - loss | 0 (0%) | 0 (0%) | 1 (1.16%) | 0 (0%) | 0 (0%) |
| | mutant - negative | 0 (0%) | 0 (0%) | 2 (2.33%) | 0 (0%) | 0 (0%) |
| | negative | 0 (0%) | 0 (0%) | 1 (1.16%) | 0 (0%) | 0 (0%) |
| | retained | 0 (0%) | 0 (0%) | 2 (2.33%) | 0 (0%) | 0 (0%) |
| | wild type | 0 (0%) | 0 (0%) | 23 (26.74%) | 0 (0%) | 13 (24.53%) |
| CDKN2A Status | unknown | 156 (100%) | 6 (100%) | 75 (87.21%) | 12 (100%) | 53 (100%) |
| | negative | 0 (0%) | 0 (0%) | 1 (1.16%) | 0 (0%) | 0 (0%) |
| | wild type | 0 (0%) | 0 (0%) | 7 (8.14%) | 0 (0%) | 0 (0%) |
| | NA | 0 (0%) | 0 (0%) | 3 (3.49%) | 0 (0%) | 0 (0%) |
| RB1 Status | unknown | 156 (100%) | 6 (100%) | 76 (88.37%) | 12 (100%) | 53 (100%) |
| | wild type | 0 (0%) | 0 (0%) | 7 (8.14%) | 0 (0%) | 0 (0%) |
| | NA | 0 (0%) | 0 (0%) | 3 (3.49%) | 0 (0%) | 0 (0%) |
| CPDEL 1p19q Status | NA | 156 (100%) | 6 (100%) | 70 (81.4%) | 12 (100%) | 20 (37.74%) |
| | mutant | 0 (0%) | 0 (0%) | 9 (10.47%) | 0 (0%) | 20 (37.74%) |
| | wild type | 0 (0%) | 0 (0%) | 7 (8.14%) | 0 (0%) | 13 (24.53%) |
| PTEN Status | NA | 156 (100%) | 6 (100%) | 78 (90.7%) | 12 (100%) | 53 (100%) |
| | mutant | 0 (0%) | 0 (0%) | 3 (3.49%) | 0 (0%) | 0 (0%) |
| | wild type | 0 (0%) | 0 (0%) | 5 (5.81%) | 0 (0%) | 0 (0%) |
| EGFR EGFRVIII Status | unknown | 156 (100%) | 6 (100%) | 73 (84.88%) | 12 (100%) | 53 (100%) |
| | EGFR Amplified | 0 (0%) | 0 (0%) | 2 (2.33%) | 0 (0%) | 0 (0%) |
| | Negative | 0 (0%) | 0 (0%) | 1 (1.16%) | 0 (0%) | 0 (0%) |
| | Positive | 0 (0%) | 0 (0%) | 1 (1.16%) | 0 (0%) | 0 (0%) |
| | wild type | 0 (0%) | 0 (0%) | 6 (6.98%) | 0 (0%) | 0 (0%) |
| | NA | 0 (0%) | 0 (0%) | 3 (3.49%) | 0 (0%) | 0 (0%) |
| Age | Mean ± SD | 11.59 ± 6.52 | 53.33 ± 9.31 | 52.29 ± 18.26 | NaN ± NA | 36.17 ± 11.7 |

[a] Children's Hospital of Philadelphia [b] City of Hope[c] University of California Los Angeles [d] University of California San Francisco [e] Glioblastoma

[f] High Grade Glioma[g] Pleomorphic Xanthroastrocytoma[h] Focal Cortical Dysplasia[i] Follicular Thyroid Cancer[j] Venous Malformation [k] follicular thyroid microcarcinoma

#### Table S2. Immune antibody panel

| Internal ID | Target | Clone | Vendor | Cat Number | Lot # | Mass | Element | Titer (µg/mL) |
|---|---|---|---|---|---|---|---|---|
| 1894 | Calprotectin | MAC387 | Thermo | MA1-80446 | WJ3419361 | 69 | Ga | 1 |
| 1893 | Mast Cell Chymase | EPR13136 | Abcam | ab233729 | GR3255393-3 | 71 | Ga | 0.25 |
| 1892 | Mast Cell Tryptase | EPR9522 | Abcam | ab271916 | GR3366012-6 | 71 | Ga | 0.25 |
| 1710 | GLUT1 | EPR3915 | Abcam | ab252403 | GR3374509-2 | 89 | Y | 0.5 |
| 1885 | CD47 | D3O7P | CST | 97253SF | 1 | 113 | In | 0.5 |
| 1771 | dsDNA | 35I9 DNA | abcam | ab27156 | GR3354725-7 | 115 | In | 1 |
| 1785 | HH3 | D1H2 | CST | 4499BF | 19 | 115 | In | 1 |
| 1902 | CD123 | IL13RA/1531 | Thermo | 3563MSM1P1 | 35631P2101 | 141 | Pr | 1 |
| 1815 | LAG3 | D2G4O | CST | 15372BF | 7 | 142 | Nd | 0.25 |
| 1872 | CD4 | EPR6855 | Abcam | ab181724 | GR3352909-7 | 143 | Nd | 0.25 |
| 1722 | ICOS | D1K2T | CST | 89601BF | 9 | 144 | Nd | 0.5 |
| 1884 | CD206 | 685645 | R&D Systems | MAB25341 | cezr0221011 | 145 | Nd | 0.5 |
| 1786 | FOXP3 | 236A/E7 | Thermo | 14-4777-37 | 2257482 | 146 | Nd | 1 |
| 1773 | PD1 | D4W2J | CST | 86163BF | 7 | 147 | Sm | 0.25 |
| 1827 | CD31 | EP3095 | abcam | ab226157 | GR3356121-6 | 148 | Nd | 0.25 |
| 1829 | PDL1 (Biotin) | 1D4-C5 | Biolegend | 409002 | B295067 | 149 | Sm | 2 |
| 1807 | ArginaseI | D4E3M | CST | 93668BF | 5 | 150 | Nd | 1 |
| 1717 | CD141 | E7Y9P | CST | 43514bf | 2 | 151 | Eu | 0.5 |
| 1850 | CD40 | EPR20735 | Abcam | ab228818 | GR3362546-5 | 152 | Sm | 1 |
| 1873 | Tox/Tox2 | E6I3Q | CST | 62886SF | 1 | 153 | Eu | 0.5 |
| 1705 | NeuN | EPR12763 | Abcam | ab209898 | GR3271481-8 | 154 | Sm | 1 |
| 1615 | NeuN | D4G40 | CST | 24307BF | 3 | 154 | Sm | 1 |
| 1790 | iNOS | SP126 | Abcam | ab238990 | GR3288157 | 155 | Gd | 2 |
| 1774 | CD68 | D4B9C | CST | 76437BF | 5 | 156 | Gd | 0.25 |
| 1813 | CD11b | EPR1344 | Abcam | ab209970 | GR3376656-2 | 157 | Gd | 0.5 |
| 1793 | CD8a | C8/144b | Thermo | 14-0085-37 | 2388408 | 158 | Gd | 0.25 |
| 1836 | CD3e | D7A6E | CST | 85061BF | 6 | 159 | Tb | 0.5 |
| 1794 | IDO | SP260 | Abcam | ab245737 | GR3286788 | 160 | Gd | 0.5 |
| 2001 | CD38 | E7Z8C | Cell signalling | 51000BF | 2 | 161 | Dy | 0.5 |
| 1901 | TIM3 | D5D5R | CST | 81229SF | 1 | 162 | Dy | 1 |
| 1566 | CD163 | D6U1J | CST | 93498BF | 2 | 163 | Dy | 1 |
| 1570 | TMEM119 | E3E4T | CST | 41134BF | 2 | 164 | Dy | 0.5 |
| 1847 | CD208 | 1010E1.01 | Dendritics | DDX0191P-100 | DDX0191-066 | 165 | Ho | 1 |
| 1900 | CD133 | D2V8Q | CST | 51917SF | 1 | 166 | Er | 2 |
| 1763 | HLADR | EPR3692 | Abcam | ab209968 | GR3371398-2 | 167 | Er | 0.5 |
| 1775 | CD14 | D7A2T | CST | 56082BF | 2 | 168 | Er | 0.5 |
| 1776 | CD45 | D9M8I | CST | 13917BF | 11 | 169 | Tm | 0.5 |
| 1877 | CD20 | L26 | Thermo | 14 0202 82 | 2340523 | 170 | Er | 0.25 |
| 1859 | Olig2 | EPR2673 | Abcam | ab220796 | gr3278799-9 | 171 | Yb | 0.5 |
| 2002 | CD86 | E2G8P | CST | 76755SF | 1 | 172 | Yb | 0.5 |
| 1848 | CD209 | DCN46 | BD Biosciences | 624087 | 1256682 | 173 | Yb | 0.5 |
| 1569 | GFAP | GA5 | Invitrogen | 14-9892-82 | 2297222 | 174 | Yb | 0.5 |
| 1828 | Ki67 | 8D5 | CST | 9449BF | 13 | 175 | Lu | 0.5 |
| 1812 | HLA 1ABC | EMR8-5 | Ionpath | 717602-100 | 20070-10(1) | 176 | Yb | 1 |

#### Table S3. Tumor antibody panel

| Internal ID | Target | Clone | Vendor | Cat Number | Lot # | Mass | Element | Titer (µg/mL) |
|---|---|---|---|---|---|---|---|---|
| 2003 | YAP TAZ | E9M8G | CST | 93622BF | 2 | 89 | Y | 1 |
| 1885 | CD47 | D3O7P | CST | 97253SF | 1 | 113 | In | 0.5 |
| 1771 | dsDNA | 35I9 DNA | abcam | ab27156 | GR3354725-7 | 115 | In | 1 |
| 1785 | HH3 | D1H2 | CST | 4499BF | 19 | 115 | In | 1 |
| 1742 | ApoE (pan) | D7I9N | CST | 13366BF | 3 | 141 | Pr | 0.5 |
| 1708 | NG2 | E3b3G | CST | 43916BF | 2 | 142 | Nd | 1 |
| 1872 | CD4 | EPR6855 | Abcam | ab181724 | GR3352909-7 | 143 | Nd | 0.25 |
| 1835 | GM2/GD2 | 1024912 | R&D Systems | MAB10503 | CNBO0120051 | 145 | Nd | 2 |
| 1786 | FOXP3 | 236A/E7 | Thermo | 14-4777-37 | 2257482 | 146 | Nd | 1 |
| 2004 | B7H3 | polyclonal | RD Systems | AF1027 | HCM0422021 | 147 | Sm | 1 |
| 1827 | CD31 | EP3095 | abcam | ab226157 | GR3356121-6 | 148 | Nd | 0.25 |
| 1703 | VISTA | D5L5T | CST | 54979BF | 2 | 149 | Sm | 0.125 |
| 1761 | EGFR | D38B1 | CST | 4267BF | 26 | 150 | Nd | 0.5 |
| 1714 | IL13RA2 | Poly | ProteinTech | 11059-1-AP | NA | 151 | Eu | 2 |
| 1825 | HER2 | 3B5 | Millipore | OP15-100ug | 3498133 | 153 | Eu | 2 |
| 1705 | NeuN | EPR12763 | Abcam | ab209898 | GR3271481-8 | 154 | Sm | 1 |
| 1615 | NeuN | D4G40 | CST | 24307BF | 3 | 154 | Sm | 1 |
| 1924 | IDH1 R132H | H09 | ARP | DIA-H09-SB-05 | 210218/05SB | 155 | Gd | 1 |
| 1846 | EGFRvIII | D6T2Q | CST | 64952BF | 2 | 156 | Gd | 0.5 |
| 1707 | EphA2 | D4A2 | CST | 6997BF | 2 | 157 | Gd | 1 |
| 1793 | CD8a | C8/144b | Thermo | 14-0085-37 | 2388408 | 158 | Gd | 0.25 |
| 1836 | CD3e | D7A6E | CST | 85061BF | 6 | 159 | Tb | 0.5 |
| 1921 | Glypican 2 | Poly | ThermoFisher | PA5-48007 | WL3458536A | 162 | Dy | 0.5 |
| 1566 | CD163 | D6U1J | CST | 93498BF | 2 | 163 | Dy | 1 |
| 1570 | TMEM119 | E3E4T | CST | 41134BF | 2 | 164 | Dy | 0.5 |
| 1716 | Histone H3.3 K27M | RM192 | RevMab | 31-1175-00 | T-11-03597 | 165 | Ho | 0.5 |
| 1900 | CD133 | D2V8Q | CST | 51917SF | 1 | 166 | Er | 2 |
| 1930 | HLADR | EPR3692 | Abcam | ab209968 | GR3371398-4 | 167 | Er | 0.5 |
| 1775 | CD14 | D7A2T | CST | 56082BF | 2 | 168 | Er | 0.5 |
| 1776 | CD45 | D9M8I | CST | 13917BF | 11 | 169 | Tm | 0.5 |
| 1738 | CD70 | poly | Lifespan Bio | A8809-50 | 126751 | 170 | Er | 0.5 |
| 1859 | Olig2 | EPR2673 | Abcam | ab220796 | gr3278799-9 | 171 | Yb | 0.5 |
| 1720 | H3K27me3 | C36B11 | CST | 9733BF | 20 | 172 | Yb | 0.25 |
| 1569 | GFAP | GA5 | Invitrogen | 14-9892-82 | 2297222 | 174 | Yb | 0.5 |
| 1828 | Ki67 | 8D5 | CST | 9449BF | 13 | 175 | Lu | 0.5 |
| 1812 | HLA 1ABC | EMR8-5 | Ionpath | 717602-100 | 20070-10(1) | 176 | Yb | 1 |

### Table S4. WHO grade classifier importance scores

| Importance | Feature Type | Feature Type Two | Features Broad | Combined Feature Type |
|---|---|---|---|---|
| 2.399543385 | MIBI | Endothelial_cells__protein_intensity__CD31 | Endothelial cell features | MIBI |
| 2.298791269 | Glycan | H6N3F1 | Fucosylated, Hybrid | Glycan |
| 2.171887082 | MIBI | GLUT1_func_counts | GLUT1_func_counts | MIBI |
| 1.350328538 | Glycan | H5N5F2 | Fucosylated | Glycan |
| 0.502899942 | RNA | VEGFA | Angiogenesis | CD45- cell RNA |
| 0.460179841 | Glycan | H5N5F3 | Fucosylated | Glycan |
| 0.299529266 | RNA | MRPS34 | MRPS34 | CD45- cell RNA |
| 0.212637889 | RNA | SCAPER | SCAPER | CD45- cell RNA |
| 0.208329012 | MIBI | Tumor_cells_Tox_over_Tumor_cells_Tox_plus_B7H3_func | Tumor antigen features | MIBI |
| 0.202173716 | RNA | CD79B | B cells | CD45- cell RNA |
| 0.192370026 | RNA | CDH2 | EMT signature | CD45- cell RNA |
| 0.153930969 | RNA | HAGH | HAGH | CD45- cell RNA |
| 0.147094602 | RNA | MEF2C | MEF2C | CD45- cell RNA |
| 0.145628655 | RNA | TM7SF3 | TM7SF3 | CD45- cell RNA |
| 0.142701038 | RNA | ABITRAM | ABITRAM | CD45- cell RNA |
| 0.120453911 | RNA | LAMTOR3 | LAMTOR3 | CD45- cell RNA |
| 0.114815128 | Glycan | H4N5F1S1 | Fucosylated, Sialylated | Glycan |
| 0.112260689 | RNA | APMAP | APMAP | CD45- cell RNA |
| 0.109882401 | RNA | TMEM44 | TMEM44 | CD45+ cell RNA |
| 0.10620754 | RNA | COL6A2 | Fibroblasts | CD45- cell RNA |
| 0.104531826 | RNA | SCAMP1 | SCAMP1 | CD45+ cell RNA |
| 0.098576938 | RNA | RAB23 | RAB23 | CD45- cell RNA |
| 0.096889452 | RNA | FN1 | Matrix | CD45- cell RNA |
| 0.096674565 | RNA | SLC43A3 | SLC43A3 | CD45- cell RNA |
| 0.095043301 | RNA | TMEM128 | TMEM128 | CD45- cell RNA |
| 0.090489337 | MIBI | Myeloid_CD11b_HLADRminus_over_all_tumor_count_prop | Immune cell features | MIBI |
| 0.089256191 | MIBI | Tumor_cells__protein_intensity__HLA1 | Tumor antigen features | MIBI |
| 0.088830115 | RNA | ATRX | ATRX | CD45+ cell RNA |
| 0.079728943 | MIBI | NA__protein_intensity__B7H3_segment_low_prop | Tumor antigen features | MIBI |
| 0.075324857 | MIBI | Myeloid_CD11b_HLADRminus_over_Unassigned_plus_Myeloid_CD11b_HLADRminus_prop | Immune cell features | MIBI |
| 0.075270371 | RNA | SOCS3 | M1 signature | CD45- cell RNA |
| 0.065517541 | RNA | TGFB2 | Protumor cytokines | CD45- cell RNA |
| 0.061789364 | MIBI | Tumor_cells__protein_intensity__B7H3 | Tumor antigen features | MIBI |
| 0.059636935 | MIBI | Immune_over_Endothelial_plus_Immune_prop | Immune_over_Endothelial_plus_Immune_prop | MIBI |
| 0.05916742 | RNA | CDK2 | Tumor proliferation rate | CD45- cell RNA |
| 0.05878816 | MIBI | Tumor_cells__protein_intensity__ApoE | Tumor antigen features | MIBI |
| 0.058387754 | MIBI | Macrophage_CD68_over_NG2_func_plus_Macrophage_CD68_prop | Tumor antigen features | MIBI |
| 0.053923645 | MIBI | NA__protein_intensity__B7H3_segment_high_prop | Tumor antigen features | MIBI |
| 0.049189668 | RNA | ARHGEF17 | ARHGEF17 | CD45- cell RNA |
| 0.048494377 | RNA | MAEA | MAEA | CD45- cell RNA |
| 0.048028843 | MIBI | B7H3_func_GPC2_func_density | Tumor antigen features | MIBI |
| 0.047570298 | RNA | CXCR4 | Myeloid cells traffic | CD45- cell RNA |
| 0.047508242 | RNA | HYAL2 | HYAL2 | CD45- cell RNA |
| 0.047465886 | RNA | RBBP6 | RBBP6 | CD45- cell RNA |
| 0.047344953 | RNA | MYG1 | MYG1 | CD45- cell RNA |
| 0.045951358 | RNA | LCAT | LCAT | CD45- cell RNA |
| 0.04114561 | Glycan | H5N2 | High Mannose | Glycan |
| 0.040899901 | RNA | PDCL3 | PDCL3 | CD45- cell RNA |
| 0.039208658 | RNA | RAB40AL | RAB40AL | CD45- cell RNA |
| 0.038864981 | RNA | CTSG | Neutrophil signature | CD45- cell RNA |
| 0.038777206 | RNA | RYK | RYK | CD45- cell RNA |
| 0.038313445 | RNA | DYSF | DYSF | CD45+ cell RNA |
| 0.037758373 | RNA | CUX1 | CUX1 | CD45+ cell RNA |
| 0.037675121 | RNA | ZNF117 | ZNF117 | CD45- cell RNA |
| 0.037481578 | MIBI | Macrophage_CD68_over_VISTA_func_plus_Macrophage_CD68_prop | Tumor antigen features | MIBI |
| 0.036745103 | RNA | KBTBD6 | KBTBD6 | CD45+ cell RNA |
| 0.036395684 | MIBI | Tumor_cells__protein_intensity__H3K27me3 | Tumor antigen features | MIBI |
| 0.035643892 | MIBI | Microglia__protein_intensity__PDL1 | Immune cell features | MIBI |
| 0.034321752 | RNA | FAM120B | FAM120B | CD45- cell RNA |
| 0.033690476 | RNA | MCM2 | Tumor proliferation rate | CD45- cell RNA |
| 0.031603463 | MIBI | Ki67_density | Ki67_density | MIBI |
| 0.031587367 | MIBI | PDL1_density | PDL1_density | MIBI |
| 0.031581699 | RNA | EEF1AKMT2 | EEF1AKMT2 | CD45- cell RNA |
| 0.031341667 | RNA | PGRMC1 | PGRMC1 | CD45- cell RNA |
| 0.031167965 | MIBI | CD31_func_counts | CD31_func_counts | MIBI |
| 0.03089169 | RNA | POLR2K | POLR2K | CD45- cell RNA |
| 0.030675365 | RNA | TXNRD1 | TXNRD1 | CD45- cell RNA |
| 0.030398182 | Glycan | H5N4F1S1_Na1 | Fucosylated, Sialylated, Biantennary | Glycan |
| 0.030297115 | MIBI | GM2_GD2_func_VISTA_func_over_all_cell_count_tumor_FOV_prop | Tumor antigen features | MIBI |
| 0.030066606 | RNA | TBC1D21 | TBC1D21 | CD45+ cell RNA |
| 0.029422222 | RNA | SEPHS1 | SEPHS1 | CD45- cell RNA |
| 0.029225361 | RNA | TBCB | TBCB | CD45+ cell RNA |
| 0.0292 | MIBI | NA__protein_intensity__GPC2_segment_high_prop | Tumor antigen features | MIBI |
| 0.02908263 | RNA | ADAMTS1 | ADAMTS1 | CD45- cell RNA |
| 0.028976608 | RNA | S100B | S100B | CD45+ cell RNA |

### Table S5. GBM survival status classifier importance scores

| Importance | Feature Type | Feature Type Two | Features Broad | Combined Feature Type |
|---|---|---|---|---|
| 0.122316799 | RNA | MCM2 | Tumor proliferation rate | CD45+ cell RNA |
| 0.105477417 | MIBI | Immune_unassigned_over_GPC2_func_plus_Immune_unassigned_prop | Tumor antigen features | MIBI |
| 0.10095592 | RNA | KIF3A | KIF3A | CD45+ cell RNA |
| 0.0998911 | RNA | THOC6 | THOC6 | CD45+ cell RNA |
| 0.091316378 | MIBI | Ki67_func_counts | Ki67_func_counts | MIBI |
| 0.085209805 | RNA | TMED3 | TMED3 | CD45+ cell RNA |
| 0.072764929 | MIBI | Tcell_CD8_Ki67_over_Tcell_CD8_Ki67_plus_VISTA_func | Tumor antigen features | MIBI |
| 0.065012987 | MIBI | Tumor_cells_Tox_over_all_cell_count_FOV_immune | Tumor antigen features | MIBI |
| 0.062836445 | RNA | EYA2 | EYA2 | CD45- cell RNA |
| 0.062438889 | MIBI | APC_over_GPC2_func_plus_APC_prop | Tumor antigen features | MIBI |
| 0.057819048 | RNA | RRAGA | RRAGA | CD45- cell RNA |
| 0.056795455 | MIBI | Immune_unassigned_over_Macrophage_CD206_plus_Immune_unassigned_prop | Immune cell features | MIBI |
| 0.054324675 | RNA | ITGB1BP1 | ITGB1BP1 | CD45+ cell RNA |
| 0.050587413 | RNA | AUH | AUH | CD45+ cell RNA |
| 0.050047543 | RNA | ZW10 | ZW10 | CD45+ cell RNA |
| 0.049786364 | RNA | TMBIM4 | TMBIM4 | CD45- cell RNA |
| 0.047459452 | RNA | CRB2 | CRB2 | CD45+ cell RNA |
| 0.044474747 | MIBI | GPC2_func_over_GPC2_func_plus_VISTA_func_prop | Tumor antigen features | MIBI |
| 0.042761364 | RNA | PDGFRB | Fibroblasts | CD45+ cell RNA |
| 0.042379293 | MIBI | Macrophage_CD206_over_Macrophage_CD68_plus_Macrophage_CD206_prop | Immune cell features | MIBI |
| 0.042099242 | MIBI | Tumor_cells_Tox_over_Tumor_cells_Tox_plus_B7H3_func | Tumor antigen features | MIBI |
| 0.041239525 | RNA | ZNF658 | ZNF658 | CD45+ cell RNA |
| 0.040721501 | RNA | GTF2H1 | GTF2H1 | CD45- cell RNA |
| 0.03964627 | RNA | ADM | ADM | CD45+ cell RNA |
| 0.039080808 | RNA | NEMF | NEMF | CD45- cell RNA |
| 0.037363636 | RNA | EDEM2 | EDEM2 | CD45+ cell RNA |
| 0.037260963 | MIBI | Tumor_cells_Tox_over_all_tumor_count | Tumor antigen features | MIBI |
| 0.03669733 | RNA | CHST7 | CHST7 | CD45- cell RNA |
| 0.036313131 | MIBI | VISTA_func_over_GPC2_func_plus_VISTA_func_prop | Tumor antigen features | MIBI |
| 0.036137626 | RNA | TMOD1 | TMOD1 | CD45- cell RNA |
| 0.035199866 | MIBI | Tumor_cells_PDL1_over_all_immune_count | Tumor antigen features | MIBI |
| 0.033481283 | RNA | SOX13 | SOX13 | CD45- cell RNA |
| 0.033084284 | RNA | CTC1 | CTC1 | CD45- cell RNA |
| 0.032977273 | MIBI | Unassigned_Ki67_over_Unassigned_Ki67_plus_VISTA_func | Tumor antigen features | MIBI |
| 0.032977273 | RNA | VPS52 | VPS52 | CD45+ cell RNA |
| 0.032878788 | MIBI | Endothelial_cells_over_Immune_unassigned_plus_Endothelial_cells_prop | Endothelial cell features | MIBI |
| 0.032788567 | RNA | CDH4 | CDH4 | CD45- cell RNA |
| 0.032244755 | RNA | SNX25 | SNX25 | CD45+ cell RNA |
| 0.030636364 | MIBI | Immune_unassigned_over_GM2_GD2_func_plus_Immune_unassigned_prop | Tumor antigen features | MIBI |
| 0.030274675 | MIBI | Endothelial_cells_over_NG2_func_plus_Endothelial_cells_prop | Endothelial cell features | MIBI |
| 0.030012987 | RNA | GPR137B | GPR137B | CD45- cell RNA |
| 0.029386364 | RNA | FCHSD2 | FCHSD2 | CD45- cell RNA |
| 0.029236364 | MIBI | Endothelial_cells_GLUT1_over_Endothelial_cells_GLUT1_plus_NG2_func | Endothelial cell features | MIBI |
| 0.028845523 | RNA | MON2 | MON2 | CD45+ cell RNA |
| 0.028159596 | MIBI | Macrophage_CD68_CD163__protein_intensity__CD68 | Immune cell features | MIBI |
| 0.026772727 | RNA | PPP1R15B | PPP1R15B | CD45- cell RNA |
| 0.025788617 | RNA | PLSCR3 | PLSCR3 | CD45+ cell RNA |
| 0.024494949 | MIBI | Macrophage_CD206_minus_Tcell_CD8 | Immune cell features | MIBI |
| 0.02425 | RNA | CALCOCO1 | CALCOCO1 | CD45- cell RNA |
| 0.024090909 | RNA | ATF7 | ATF7 | CD45+ cell RNA |
| 0.024060829 | RNA | TCTA | TCTA | CD45- cell RNA |
| 0.023801768 | RNA | TIMM23 | TIMM23 | CD45+ cell RNA |
| 0.023427273 | MIBI | Macrophage_CD206_over_VISTA_func_plus_Macrophage_CD206_prop | Tumor antigen features | MIBI |
| 0.023345455 | MIBI | Immune_unassigned_over_Unassigned_plus_Immune_unassigned_prop | Immune cell features | MIBI |
| 0.022922727 | MIBI | Immune_unassigned_over_Endothelial_cells_plus_Immune_unassigned_prop | Endothelial cell features | MIBI |
| 0.022636364 | RNA | COX7A1 | COX7A1 | CD45- cell RNA |
| 0.022495637 | MIBI | Myeloid_CD14__protein_intensity__Ki67 | Immune cell features | MIBI |
| 0.022263636 | MIBI | Immune_unassigned_over_HER2_func_plus_Immune_unassigned_prop | Tumor antigen features | MIBI |
| 0.021969697 | RNA | ZNF608 | ZNF608 | CD45- cell RNA |
| 0.021909091 | RNA | MBD4 | MBD4 | CD45- cell RNA |
| 0.021745455 | RNA | NOP53 | NOP53 | CD45- cell RNA |
| 0.021507081 | RNA | SPG11 | SPG11 | CD45- cell RNA |
| 0.021272727 | MIBI | Immune_unassigned_over_Tcell_CD4_plus_Immune_unassigned_prop | Immune cell features | MIBI |
| 0.021090909 | MIBI | Immune_unassigned_over_Microglia_CD163_plus_Immune_unassigned_prop | Immune cell features | MIBI |
| 0.020181818 | RNA | RBBP9 | RBBP9 | CD45- cell RNA |
| 0.020084632 | MIBI | Tox_func_counts | Tox_func_counts | MIBI |
| 0.019727273 | MIBI | Endothelial_over_NG2_func_plus_Endothelial_prop | Tumor antigen features | MIBI |
| 0.019363636 | MIBI | Tox_func_counts | Tox_func_counts | MIBI |
| 0.018990404 | RNA | GNG7 | GNG7 | CD45- cell RNA |
| 0.018902597 | RNA | GNE | GNE | CD45+ cell RNA |
| 0.018440794 | MIBI | Macrophage_CD68_over_Macrophage_CD206_plus_Macrophage_CD68_prop | Immune cell features | MIBI |
| 0.017949495 | MIBI | Tox_func_Ki67_func_counts | Tox_func_Ki67_func_counts | MIBI |
| 0.01779021 | RNA | SIPA1L3 | SIPA1L3 | CD45- cell RNA |
| 0.017466977 | RNA | GRIN2D | GRIN2D | CD45- cell RNA |
| 0.017454545 | RNA | ZNF419 | ZNF419 | CD45- cell RNA |
